# Localized Rigidification and Allosteric Modulation Mechanisms of SARS-CoV-2 Spike Neutralization by Class 3 and Class 4 Antibodies at Atomic Resolution: An Integrated Computational Study of Binding, Dynamics, and Allostery

**DOI:** 10.64898/2026.07.12.738055

**Authors:** Mohammed Alshahrani, Will Gatlin, Max Ludwick, Lucas Turano, Brandon Foley, Gennady Verkhivker

**Affiliations:** Keck Center for Science and Engineering, Department of Biological Sciences, Schmid College of Science and Technology, Chapman University, Orange, CA 92866, United States of America; Department of Biomedical and Pharmaceutical Sciences, Chapman University School of Pharmacy, Irvine, CA 92618, United States of America; Department of Pharmacology, Skaggs School of Pharmacy and Pharmaceutical Sciences, University of California San Diego, 9500 Gilman Drive, La Jolla, CA 92093, United States of America

## Abstract

The relentless evolution of SARS-CoV-2 and the emergence of highly antibody-evasive variants underscore the need to decipher the molecular principles that govern antibody neutralization breadth and resilience. In this study, we employ an integrated computational framework combining structural analysis, conformational dynamics, mutational scanning, binding energetics, and allosteric network modeling to dissect the mechanistic signatures of class 3 and class 4 antibodies targeting the receptor-binding domain (RBD) of the SARS-CoV-2 spike protein. Through comprehensive analysis of antibody-RBD complexes including individual antibodies (COV2-3835, COV2-3891, COV2-3906) and synergistic dual-antibody pairs we uncover a fundamental mechanistic dichotomy that distinguishes these two antibody classes and explains their differential patterns of neutralization potency, breadth, and resilience to viral escape. Our analysis reveals that class 3 antibodies achieve neutralization with mechanical perturbation strictly confined to the binding interface. In contrast, class 4 antibodies employ a long-range allosteric destabilization mechanism, anchoring to a structurally rigid hydrophobic core and establishing a mechanical conduit through the β-sheet core that transmits conformational changes. Mutational scanning and rigorous energetic analysis reveal fundamentally different vulnerability landscapes: class 4 epitopes are defined by an immutable hydrophobic core that is exquisitely sensitive to mutation yet evolutionarily constrained across sarbecoviruses, explaining their ultra-broad binding and limited escape potential. Class 3 epitopes exhibit a plastic periphery with a conserved anchor and variable sensitivity in peripheral regions, creating multiple escape pathways. These predictions show excellent agreement with experimental deep mutational scanning data, validating our computational approach and establishing a quantitative framework for predicting immune escape. Allosteric network analysis identifies the β-sheet core as the critical communication conduit for class 4 antibodies, with specific residues serving as essential hubs that connect the hydrophobic core to the RBM loop. The convergence of high communication centrality with extreme perturbation sensitivity at these positions establishes them as the most critical allosteric hotspots, essential for function and resistant to mutation. The proposed multi-pronged computational framework provides a generalizable approach for understanding antibody neutralization mechanisms and predicting immune escape across diverse viral targets, with implications for the rational design of next-generation antibody therapeutics that balance potency, breadth, and resilience.

## Introduction

The emergence of severe acute respiratory syndrome coronavirus 2 (SARS-CoV-2) has prompted an intensive global research effort to understand its molecular architecture, mechanisms of host cell entry, and the immune responses it elicits. Central to these investigations is the Spike (S) glycoprotein, a trimeric surface structure that mediates viral entry and serves as the primary target of neutralizing antibodies.^1–15^ The S protein exhibits remarkable conformational flexibility, transitioning through multiple functional states - from receptor engagement to membrane fusion while simultaneously evading immune surveillance. Comprising two distinct subunits, S1 and S2, the S protein includes within S1 the N-terminal domain (NTD), the receptor-binding domain (RBD), and two conserved subdomains (SD1, SD2) that stabilize the prefusion conformation.^16–18^ The RBD, in particular, plays a critical role in binding the angiotensin-converting enzyme 2 (ACE2) receptor, making it a focal point for neutralizing antibody responses. Extensive cryo-electron microscopy (cryo-EM) and X-ray structures of SARS-CoV-2 variants of concern (VOCs) in various functional states, along with their interactions with antibodies, underscore how VOCs induce structural changes in the dynamic equilibrium of the S protein.^19–25^ These findings reflect the balance between structural stability, immune evasion, and receptor binding that shapes the evolutionary trajectory of SARS-CoV-2 and its variants.

The continued evolution of SARS-CoV-2 within the Omicron lineage and its descendants, including XBB.1, XBB.1.5, JN.1, KP.2, KP.3, and more recent subvariants, highlights the extraordinary virus adaptability and the emergence of convergent evolutionary hotspots: specific residues repeatedly and independently mutated across geographically and temporally distinct lineages.^26–29^ The recurrence of mutations such as F456L, R346T, L455F, and K444T signals a narrowing evolutionary landscape where SARS-CoV-2 optimizes a limited set of high-impact changes to balance transmissibility, immune evasion, and structural stability. The JN.1-derived subvariants KP.2 and KP.3 have independently acquired constellations of key S mutations, including R346T, F456L, Q493E, and V1104L, that collectively enhance both transmissibility and the ability to evade neutralizing antibodies.^26–29^ Recent cryo-EM structural analyses reveal that the F456L mutation synergizes with Q493E to strengthen ACE2 binding, providing a structural basis for enhanced infectivity and immune resistance in lineages such as KP.3.^30,31^ The emergence of multiple antibody-evasive variants KP.3.1.1, XEC, LP.8.1, LF.7, NB.1.8.1, XFG, and BA.3.2 reflects continued adaptation under immune selection pressure, with the evolutionary trajectory transitioning from discrete point mutations toward complex multilineage recombination events that enhance transmissibility and immune escape.^32–39^ Together, these variants illustrate that SARS-CoV-2 evolution has increasingly narrowed to optimize a limited set of high-impact changes, underscoring the urgent need to identify and characterize antibodies targeting epitopes where mutational escape imposes prohibitive fitness costs.

The immune response to SARS-CoV-2 is characterized by a remarkable diversity of antibodies that target distinct regions of the S protein, with the RBD serving as the primary focus of neutralizing activity.^40–48^ This diversity reflects the ability of the mammalian immune system to recognize virtually every solvent-accessible residue on the RBD surface, generating antibodies with varying epitope specificities, binding geometries, and neutralization mechanisms.^48^ The structural characterization of antibody-RBD complexes is fundamental to understanding the molecular mechanisms underlying neutralization of SARS-CoV-2. Antibodies targeting the RBD can be classified according to several complementary systems that capture different aspects of their recognition properties. Structural classifications group antibodies based on their epitope location on the RBD and their preference for “up” or “down” RBD conformations, while functional classifications derived from deep mutational scanning (DMS) partition antibodies according to their escape mutation profiles and neutralization breadth.^40–45^ These classification frameworks have proven essential for understanding the molecular determinants of antibody potency, the patterns of viral escape, and the identification of antibodies with exceptional resilience to antigenic drift. The Barnes classification divides SARS-CoV-2 RBD-targeting antibodies into four structural classes based on epitope location and RBD conformation.^40^ Class 1 antibodies recognize “up” RBD conformations, overlapping with the ACE2-binding motif; class 2 antibodies bind “down” RBD conformations, also targeting the ACE2-binding region; class 3 antibodies engage the side of the RBD away from the receptor-binding motif; and class 4 antibodies target cryptic or conserved sites on the RBD underside, accessible only in the “up” conformation. Class 1 and 2 antibodies are the most potent and frequently elicited, targeting overlapping regions of the RBD where ACE2 binds generally achieving high potency through direct competition with ACE2 but are readily escaped by single amino acid substitutions, whereas antibodies targeting more conserved epitopes outside the RBM often exhibit broader cross-reactivity but lower potency due to indirect mechanisms of neutralization.^40,41^ In contrast, class 3 and 4 antibodies are less potent and less frequently elicited in humans but target relatively more conserved RBD regions and exhibit cross-reactivity with VOCs.^40,41^ High-throughput yeast display and DMS have enabled systematic mapping of antibody epitopes and escape mutations, leading to comprehensive classification of neutralizing antibodies into six major epitope groups (A–F) based on their binding footprints.^42,43^ The six-group classification, derived from large-scale DMS profiling, captures nuanced differences in escape patterns where groups A-D correspond to Barnes classes 1 and 2, subdivided based on sensitivity to specific mutations; group E maps to Barnes class 3, targeting the RBD side with relative resilience to RBM mutations; and group F corresponds to Barnes class 4, targeting conserved cryptic epitopes and demonstrating broad neutralization.^42–45^ Beyond the Barnes structural framework, multiple complementary classification systems have been developed to capture the complexity of antibody-RBD recognition. The RBD-1 to RBD-7 scheme provides enhanced resolution while preserving the overarching hierarchical organization, with RBD-1/2 mapping to Barnes classes 1/2 and Cao groups A-D, RBD-3/4/5 corresponding to Barnes class 3 and Cao group E, and RBD-6/7 aligning with Barnes class 4 and Cao group F antibodies.^46^ Structure-based approaches have further refined this landscape by employing quantitative residue contact definitions and clustering methodologies, identifying 23 frequently engaged epitope sites and characterizing the topological features of RBD surfaces that mediate antibody and nanobody interactions.^47^ Large-scale systematic analyses of antibody recognition have categorized RBD-targeting antibodies into four epitope groups based on residue overlap, with groups I and II directed toward the highly plastic receptor-binding site and groups III and IV engaging more evolutionarily stable non-RBS regions.^47,48^ These integrated analyses have revealed that the antibody repertoire can access nearly the entire RBD solvent-accessible surface, with approximately 99% coverage, underscoring the remarkable capacity of the humoral immune system to recognize diverse epitope landscapes.^48^

The discovery of antibodies capable of maintaining efficacy across highly divergent viral lineages has illuminated distinct mechanistic strategies for achieving broad neutralization. Class 1 antibodies including BD55-1205,^45^ VIR-7229,^49^ 19-77,^50^ ZCP3B4, and ZCP4C9^51^ each employ unique molecular approaches to achieve cross-variant activity. While the majority of clinically approved monoclonal antibodies have lost effectiveness against Omicron sublineages and emerging JN.1 derivatives class 4 antibodies of the F3 subgroup pemivibart (VYD222)^52^ and SA55 (BD55-5514)^53^ have demonstrated exceptional neutralization potency against the most recent JN.1 descendant variants. The persistent emergence of antibody-evasive variants has highlighted the urgent need to identify and characterize antibodies directed against conserved epitopes, where mutational escape entails substantial fitness penalties. A pivotal advance in this pursuit was achieved by Cao and colleagues, who systematically interrogated orphan broadly reactive RBD-binding antibodies and uncovered three conserved epitope sites that remain accessible throughout SARS-CoV-2 evolution.^54^ Another newly characterized highly conserved region on the RBD targeted by a special class of broadly neutralizing antibodies that resist extreme antigenic drift across variants include CC25.4, CC25.17, and CC25.56 antibodies which demonstrate broad neutralization with less potency than group 1 antibodies but significantly greater escape resistance.^55^

The growing interest to class 3 and 4 antibodies has led to emergence of new groups of antibodies from these classes that enhanced our understanding of previously unappreciated diversity of antibody engagements for these classes targeting relatively conserved RBD regions. The first group consists of dual-antibody complexes which represent cryo-EM structures of the SARS-CoV-2 S protein bound to pairs of non-overlapping neutralizing antibodies that achieve broad, synergistic neutralization. These structures are particularly informative because they reveal how two antibodies invariably including at least one class 4 antibody (such ads 3E2 or 1C4) can engage distinct epitopes simultaneously to enhance neutralization through cooperative allosteric effects.^56^ Of special interest is the complex featuring two class 4 antibodies 3E2 and 1C4 binding simultaneously to the RBD without a class 3 partner antibody that directly engages the RBM loop.^56^ This complex provides a remarkable opportunity to understand how two antibodies from the same class can achieve synergistic neutralization through complementary engagement of distinct regions on the inner face, and what this reveals about the allosteric mechanism of class 4 antibodies. The second group consists of single-antibody complexes including class 3 COV2-3835 and COV2-3891 antibodies targeting the lateral face, and class 4 COV2-3906 4 antibody targeting the cryptic inner face, providing detailed architectural information about their respective epitopes.^57^ The class 3 antibody family exhibits significant structural diversity, with binding modes ranging from extensive to compact footprints. COV2-3835 engages the lateral face through a broad epitope s while COV2-3891 exhibits a more compact footprint.^57^ This variation in epitope extent correlates with differences in conformational constraint and neutralization potency, while the class 4 antibody family exhibits remarkable structural conservation. Together, these structures reveal the epitope architectures that underlie the distinct neutralization mechanisms of class 3 and class 4 antibodies and provide the structural foundation for interpreting their conformational dynamics and allosteric communication.

The interplay between structural dynamics and binding energetics is central to understanding how antibodies achieve broad neutralization and how viruses evolve to escape them. However, a detailed understanding of dynamics and energetics underlying the unique neutralizing capacity of these novel class 3 and class 4 antibodies is lacking. The balance of structural stability and adaptability when targeting conserved epitopes is an important driver of neutralization antibody efficiency, and quantifying molecular determinants and specific hotspots of immune escape-resistant targets is of significant interest. In the current study, we address these questions from a unified biophysical perspective by integrating structure, dynamics, energetics, and allosteric network analyses to investigate how these class 3 and class 4 antibodies engage the RBD and achieve their distinct neutralization mechanisms.

Conformational dynamics and allosteric interactions can be linked to binding of novel human antibodies^58,59^ where antibody-induced associated changes in S dynamics can distinguish weak, moderate, and strong neutralizing antibodies.^60^ Recent computational and structural studies suggest that the pattern of specific escape mutants for ultrapotent antibodies may be driven by a complex balance between the impact of mutations on structural stability, binding strength, and long-range communications.^61–64^ Our recent computational studies examined ultrapotent and broadly neutralizing class 1 and class 4/1 antibodies to reveal a unifying biophysical principle that explains how these antibodies achieve both exceptional potency and remarkable resilience against viral evolution.^65,66^ These studies found that neutralizing antibodies bind via rigid, pre-configured interfaces that distribute binding energy across a broad epitope through numerous suboptimal, yet synergistic, interactions. This proposed “distributed redundancy” architecture reconciles an apparent paradox: how can ultrapotent antibodies rely on “suboptimal” contacts and still neutralize effectively? The answer lies not in maximizing affinity at a few residues, but in improving robustness through redundancy, backbone engagement, and strategic targeting of evolutionarily constrained regions.

Here, we further expand on these hypotheses by using multi-pronged computational approach that integrates conformational dynamics of antibody-RBD complexes, systematic mutational scanning to identify binding hotspots and predict escape mutations, MM-GBSA binding free energy calculations with residue-based decomposition, and allosteric network analysis to examine adaptive evolution. Our analysis focuses on several key questions: (a) what structural and energetic features distinguish class 3 and class 4 antibody binding mechanisms and neutralization strategies? (b) how do these antibody classes differ in their conformational dynamics and allosteric communication networks? (c) what are the mutational vulnerabilities and escape pathways for each antibody class, and what fitness costs might such mutations incur? (d) how do class 3 and class 4 antibodies compare in terms of neutralization potency, breadth, and resilience to viral escape?

Our results reveal that class 3 antibodies exert neutralization primarily through direct steric blockade of ACE2 engagement, with localized rigidification of the binding interface, while class 4 antibodies exert their effects primarily through long-range allosteric destabilization of the RBM. The mechanical analysis reveals a fundamental paradox of class 4 antibodies: they achieve ultra-broad binding through anchoring to a conserved, mechanically rigid core, yet exhibit weak neutralization because the allosteric mechanism—while effective at loosening the RBM—is inherently less efficient than direct steric blockade. The extreme core rigidity ensures stable binding to a conserved epitope that is resistant to mutation, explaining the ultra-broad binding of class 4 antibodies. However, the dramatic RBM loosening impairs ACE2 engagement through an indirect mechanism that relies on long-range mechanical communication, which cannot achieve the same level of potency as direct receptor blockade. Together, these insights provide a roadmap for developing next-generation antibody therapeutics that balance potency, breadth, and resilience against viral evolution.

## Materials and Methods

### Coarse-Grained Simulations

The crystal and cryo-EM structures of the RBD-antibody are obtained from the Protein Data Bank.^67^ We employed CABS-flex approach that efficiently combines a high-resolution coarse-grained (CG) model and efficient search protocol capable of accurately reproducing all-atom MD simulation trajectories and dynamic profiles of large biomolecules on a long time scale.^68–73^ In this high-resolution model, the amino acid residues are represented by Cα, Cβ, the center of mass of side chains and another pseudoatom placed in the center of the Cα-Cα pseudo-bond. In this model, the amino acid residues are represented by Cα, Cβ, the center of mass of side chains and the center of the Cα-Cα pseudo-bond. The CABS-flex approach implemented as a Python 2.7 object-oriented standalone package was used in this study to allow for robust conformational sampling proven to accurately recapitulate all-atom MD simulation trajectories of proteins on a long time scale. Conformational sampling in the CABS-flex approach is conducted with the aid of Monte Carlo replica-exchange dynamics and involves local moves of individual amino acids in the protein structure and global moves of small fragments.

The default settings were used in which soft restraints are imposed only on pairs of residues fulfilling the following conditions: the distance between their *C*α atoms was smaller than 8 Å, and both residues belong to the same secondary structure elements. A total of 1000 independent CG-CABS simulations were performed for each of the systems studied. In each simulation, the total number of cycles was set to 10,000 and the number of cycles between trajectory frames was 100.

### All-Atom Molecular Dynamics Simulations

Structural analysis and all-atom MD simulations were performed according to the protocol detailed in our recent studies of RBD-antibody complexes.^74^ In brief, the protonation states for all the titratable residues of the antibody and RBD proteins were assigned at pH 7.0 using Propka 3.1 software and web server.^75,76^ The glycan chains were built using CHARMM-GUI Glycan Reader^77,78^ at glycosylation sites N331 and N343 of RBD. NAMD 2.13-multicore-CUDA package^79^ with CHARMM36m force field^80^ were used in all-atom MD simulations. These simulations incorporate a minimal glycan representation at key structural sites, providing a realistic assessment of steric effects and surface accessibility without the prohibitive cost of modeling full glycan ensembles. This approach ensures that the local steric footprint and chemical environment of the RBD are accurately represented, particularly in regions where glycans may modulate antibody binding or receptor interaction. Each system was solvated with TIP3P water molecules and neutralizing 0.15 M NaCl in a periodic box that extended 10 Å beyond any protein atom in the system.^81^ The heavy atoms in the complex were restrained using a force constant of 1000 kJ mol^−1^ nm^−1^ to perform 1ns equilibration simulation. Long-range, non-bonded van der Waals interactions were computed using an atom-based cutoff of 12 Å, with the switching function beginning at 10 Å and reaching zero at 14 Å. The SHAKE method was used to constrain all the bonds associated with hydrogen atoms. The simulations were run using a leap-frog integrator with a 2 fs integration time step. The ShakeH algorithm in NAMD was applied for the water molecule constraints. A 310 K temperature was maintained using the Nóse-Hoover thermostat with 1.0 ps time constant and 1 atm pressure was maintained using isotropic coupling to the Parrinello-Rahman barostat.^82,83^ The long-range electrostatic interactions were calculated using the particle mesh Ewald method^84^ with a cut-off of 1.2 nm and a fourth-order (cubic) interpolation. The simulations were performed under an NPT ensemble with a Langevin thermostat and a Nosé–Hoover Langevin piston at 310 K and 1 atm. The damping coefficient (gamma) of the Langevin thermostat was 1/ps. In NAMD, the Nosé–Hoover Langevin piston method is a combination of the Nosé–Hoover constant pressure method^85,86^ and piston fluctuation control implemented using Langevin dynamics.^87^ An NPT production simulation was run on equilibrated structures for 500 ns keeping the temperature at 310 K and a constant pressure (1 atm).

### Distance Fluctuations Analysis of Protein Stability and Allosteric Propensities

Using a protein mechanics-based approach^88–91^ we probed residue stability and communication propensities with the aid of distance fluctuation analysis of the simulation trajectories for the studied SARS-CoV-2 S proteins. The fluctuations of the mean distance between a given residue and all other residues in the ensemble were converted into distance fluctuation stability and communication indexes that measure the energy cost of the residue deformation during simulations. Our previous studies^92,93^ and related models^94,95^ adapted this metric to characterize allosteric communication propensities of protein residues through their mechanical properties since mean square deformations of a given residue with respect to the rest of the protein are related to the inter-residue communication strength.

We computed the fluctuations of the mean distance between each atom within a given residue and the atoms that belong to the remaining residues of the protein. The high values of distance fluctuation indexes are associated with residues that display small fluctuations in their distances to all other residues, while small values of this stability parameter would point to more flexible sites that experience large deviations of their inter-residue distances. In our model, the distance fluctuation stability index for each residue is calculated by averaging the distances between the residues over the simulation trajectory using the following expression:

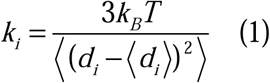

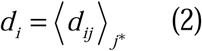

*d_ij_* is the instantaneous distance between residue *i* and residue *j*, *k_B_* is the Boltzmann constant, *T* =300K. 〈〉 denotes an average taken over the MD simulation trajectory and *d_i_* = 〈*d_ij_*〉*_j_* is the average distance from residue *i* to all other atoms *j* in the protein (the sum over *j*_*_ implies the exclusion of the atoms that belong to the residue *i*). The interactions between the *C*_α_ atom of residue *i* and the *C*_α_ atom of the neighboring residues *i*-1 and *i*+1 are excluded in the calculation since the corresponding distances are nearly constant. The inverse of these fluctuations yields an effective force constant *k_i_* that describes the ease of moving an atom with respect to the protein structure.

### Mutational Scanning Profiling

Mutational scanning analysis of the binding epitope residues for the S RBD–antibody complexes. Each binding epitope residue was systematically mutated using all substitutions, and corresponding protein stability and binding free energy changes were computed. The BeAtMuSiC approach^96–98^ was employed and evaluated the impact of mutations on both the strength of interactions at the protein–protein interface and the overall stability of the complex using statistical energy functions. BeAtMuSiC is a knowledge-based statistical potential was applied to 1,000 atomistic reconstructed conformations sampled from each CG-CABS ensemble. This method quantifies mutation-induced energy changes (ΔΔG) through three physically grounded terms: (a) a Lennard-Jones potential for van der Waals interactions parameterized on observed atomic contact frequencies in the PDB; (b) a directional hydrogen-bonding term based on geometric and distance constraints; and (c) a solvation term modeling changes in buried surface area using statistical potentials. BeAtMuSiC identifies a residue as part of the protein–protein interface if its solvent accessibility in the complex is at least 5% lower than its solvent accessibility in the individual protein partner(s). The binding free energy of the protein–protein complex can be expressed as the difference in the folding free energy of the complex and folding free energies of the two protein binding partners. The change in the binding energy due to a mutation is:

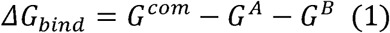

*G^com^* is the free energy of the complex. This is the Gibbs free energy associated with the folded, bound state of the entire protein–protein complex (e.g., the Spike RBD–antibody complex). *G^A^* is the free energy of the first binding partner (e.g., the isolated S-RBD) in its unbound, folded state. is the free energy of the second binding partner (e.g., the isolated antibody) in its unbound, folded state. The change in the binding energy due to a mutation was calculated then as

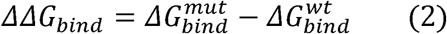

*ΔΔG_bind_* is the change in binding free energy resulting from a specific mutation. This quantifies how the mutation affects the binding affinity compared to the wild-type (original) interaction. A positive *ΔΔG_bind* typically indicates weakened binding (the mutation makes binding less favorable or more difficult), while a negative *ΔΔG_bind* indicates strengthened binding (the mutation makes binding more favorable). 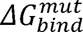 is the binding free energy calculated using Equation (4), but for the mutated protein complex (e.g., a mutant RBD bound to the antibody). 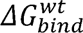 is the binding free energy calculated using Equation (4), but for the wild-type (unmutated) protein complex, serving as the reference state. We leveraged rapid calculations based on statistical potentials to compute the ensemble-averaged binding free energy changes using equilibrium samples from simulation trajectories. The binding free energy changes were obtained by averaging over 1000 and 10,000 equilibrium samples for each of the systems studied.

### Binding Free Energy Computations of the RBD Complexes with Antibodies

MM-GBSA approach^99–105^ with the AMBER21 suite^106^ was employed for rigorous validation and residue-level decomposition. This dual framework leverages BeAtMuSiC efficiency for rapid mutational screening and physical rigor of MM-GBSA computations for mechanistic dissection of van der Waals and electrostatic contributions to binding and identification of binding affinity hotspots. We calculated the ensemble-averaged changes in binding free energy using 1000 equilibrium samples obtained from simulation trajectories for each system under study. Initially, the binding free energies of the RBD–antibody complexes were assessed using the MM-GBSA approach. Additionally, we conducted an energy decomposition analysis to evaluate the contribution of each amino acid during the binding of the RBD to antibodies. The binding free energy for the RBD–antibody complex was obtained using:

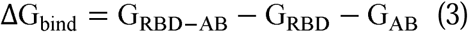

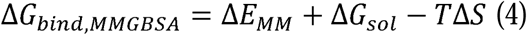

where ΔE_MM_ is total gas phase energy (sum of ΔEinternal, ΔEelectrostatic, and ΔEvdw); ΔGsol is sum of polar (ΔGGB) and non-polar (ΔGSA) contributions to solvation. Here, G _RBD–ANTIBODY_ represent the average over the snapshots of a single trajectory of the complex, G_RBD_ and G_ANTIBODY_ corresponds to the free energy of RBD and antibody, respectively.

The polar and non-polar contributions to the solvation free energy were calculated using a Generalized Born solvent model and consideration of the solvent-accessible surface area. MM-GBSA was employed to predict the binding free energy and decompose the free energy contributions to the binding free energy of a protein–protein complex on a per residue basis. The binding free energy with MM-GBSA was computed by averaging the results of computations over 10,000 samples from the equilibrium ensembles. The standard error of the mean (SEM) for binding free energy estimates was calculated from the distribution of values obtained across the 10,000 snapshots sampled for each system. In this study, we chose the “single trajectory” protocol (one trajectory of the complex) because it is less noisy due to the cancellation of intermolecular energy contributions. Entropy calculations typically dominate the computational cost of MM-GBSA estimates. In this study, the entropy contribution was not included in the calculations of binding free energies of the RBD–antibody complexes because the entropic differences in estimates of relative binding affinities were expected to be small owing to the small mutational changes and the preservation of the conformational dynamics. MM-GBSA energies were evaluated with the MMPBSA.py script in the AmberTools21 package^107^ and gmx_MMPBSA, a new tool to perform end-state free energy calculations from CHARMM and GROMACS trajectories.^108^

### Allosteric Interactions Modeling Using Residue Interaction Networks

To analyze protein structures, we employed a graph-based representation where residues are modeled as network nodes, and non-covalent interactions between residue side-chains define the edges.^109,110^ This approach captures the spatial and functional relationships between residues, providing insights into the protein structural and dynamic properties.

Residue Interaction Network Generator (RING) program^111–113^ was employed for generation of the residue interaction networks using the conformational ensemble where edges have an associated weight reflecting the frequency in which the interaction present in the conformational ensemble. The residue interaction network files in xml format were obtained for all structures using RING v3.0 webserver.^111–113^ Network graph calculations were performed using the python package NetworkX.^114^ Using the constructed protein structure networks, we computed the residue-based short path betweenness parameter. The short path betweenness of residue *i* is defined to be the sum of the fraction of shortest paths between all pairs of residues that pass through residue *i*:

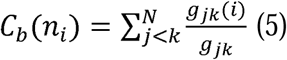

where *g_jk_* denotes the number of shortest geodesics paths connecting *j* and *k,* and *g_jk_* (*i*) is the number of shortest paths between residues *j* and *k* passing through the node *n_i_*. Residues with high occurrence in the shortest paths connecting all residue pairs have a higher betweenness values. For each node *n,* the betweenness value is normalized by the number of node pairs excluding *n* given as(*N* −1)(*N* - 2) / 2, where *N* is the total number of nodes in the connected component that node *n* belongs to. The normalized short path betweenness of residue *i* can be expressed as:

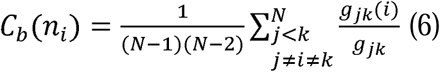

*g_jk_* is the number of shortest paths between residues *j* and k; *g_jk_* (*i*) is the fraction of these shortest paths that pass through residue *i*.

Through mutation-based perturbations of protein residues we compute dynamic couplings of residues and changes in the short path betweenness centrality (SPC) averaged over all possible modifications in a given position.

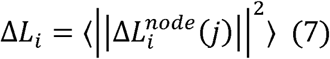

where *i* is a given site, *j* is a mutation and 〈…〉 denotes averaging over mutations. 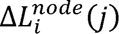 describes the change of SPC parameters upon mutation *j* in a residue node *i*. Δ*L_i_* is the average change of ASPL triggered by mutational changes in position *i*. Z-score is then calculated for each node as follows^115,115^:

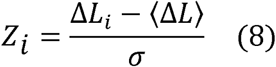

〈Δ*L*〉 is the change of the SPC network parameter under mutational scanning averaged over all protein residues and σ is the corresponding standard deviation. The ensemble-average Z score changes are computed from network analysis of the conformational ensembles of antibody-RBD complexes using 1,000 snapshots of the simulation trajectory.

## Results

### Structural Characterization of RBD Epitopes in Class 3 and Class 4 Antibody Complexes

To establish the structural basis for understanding the distinct neutralization mechanisms of class 3 and class 4 antibodies, we analyzed a comprehensive panel of antibody-RBD complexes comprising two complementary groups of structures (Figure 1). The first group comprises single-antibody complexes of COV2-3835 (Figure 1A), COV2-3891 (Figure 1B), and COV2-3906 (Figure 1C), which represent high-resolution crystal structures of individual COV2 series antibodies bound to the RBD. These structures confirmed that COV2-3835 and COV2-3891 are class 3 antibodies targeting the lateral face, while COV2-3906 is a class 4 antibody targeting the cryptic inner face, providing detailed architectural information about their respective epitopes.

**Figure 1.**
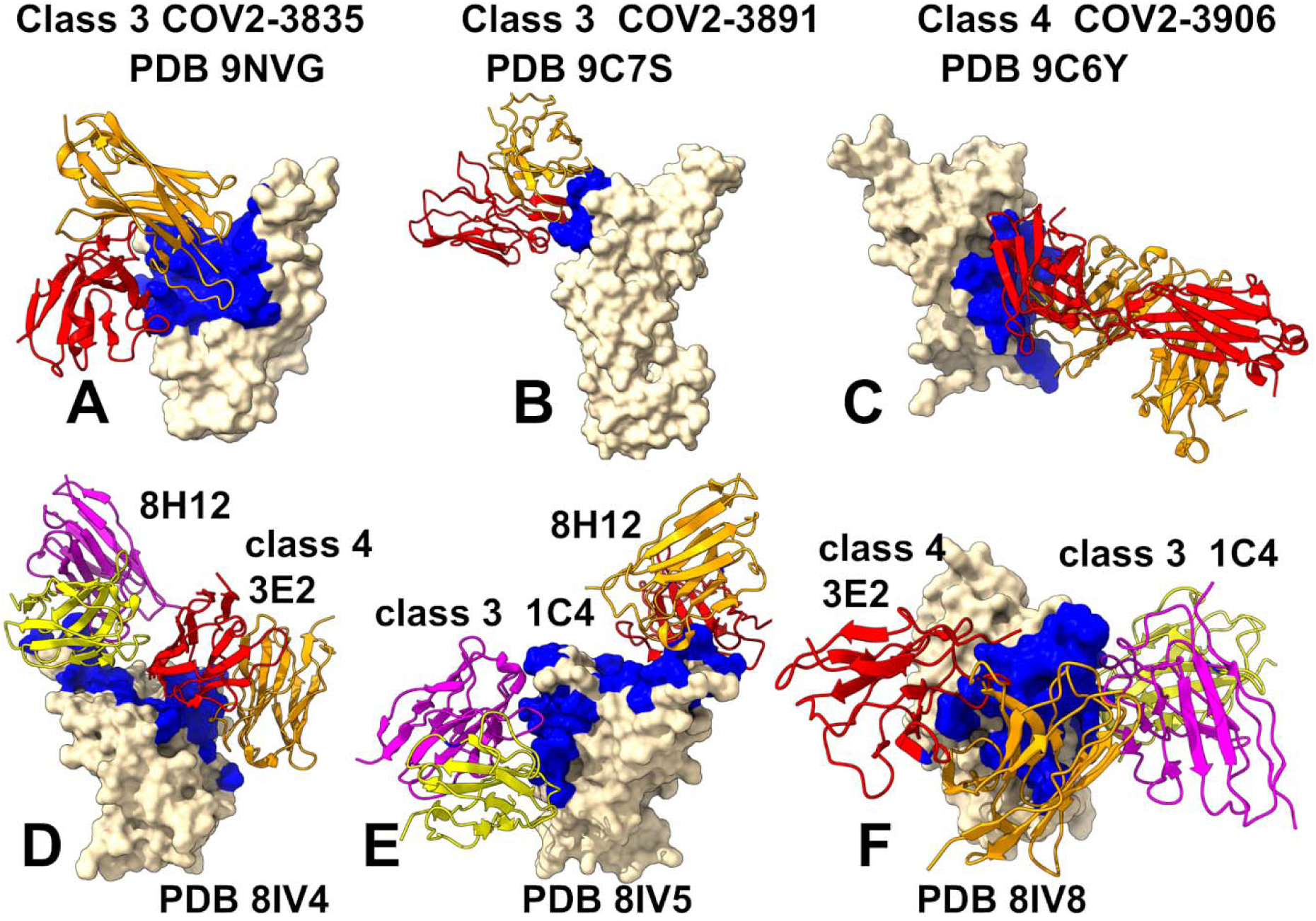
Structural architecture of class 3 and class 4 antibody-RBD complexes. (A) Class 3 antibody COV2-3835 (PDB 9NVG) bound to the RBD. The RBD is shown in surface representation (wheat color) with antibodies depicted as cartoons. The binding epitope residues are highlighted in blue surface. Heavy chain is in orange cartoon, light chain in red cartoon. (B) Class 3 antibody COV2-3891 (PDB 9C7S) bound to the RBD. Heavy chain is in orange cartoon, light chain in red cartoon. (C) Class 4 antibody COV2-3906 (PDB 9C6Y) bound to the RBD. Heavy chain is in orange cartoon, light chain in red cartoon. (D) Dual complex of class 1 antibody 8H12 and class 4 antibody 3E2 (PDB 8IV4) bound to the RBD, revealing focused 3E2 engagement of the hydrophobic core coupled with 8H12-mediated ACE2-competitive constraint on the RBM loop. 8H12 heavy and light chains are in magenta and yellow, 3E2 heavy and light chains are on orange and red. (E) Dual complex of class 1 antibody 8H12 and class 3 antibody 1C4 (PDB 8IV5), showing 1C4-mediated lateral face engagement and dual steric constraint on the RBM loop. 1C4 heavy and light chains are in magenta and yellow, 8H12 heavy and light chains are on orange and red. (F) Dual complex of class 3 antibody 1C4 and class 4 antibody 3E2 (PDB 8IV8), revealing cooperative engagement of both the lateral face (1C4) and cryptic inner face (3E2), forming an almost contiguous joined footprint covering conserved RBD regions. 3E2 heavy and light chains are on orange and red, 1C4 heavy and light chains are in magenta and yellow.

The second group comprises cryo-EM structures of the S protein bound to pairs of antibodies that achieve broad, synergistic neutralization: class 1 antibody 8H12 paired with class 4 antibody 3E2 (Figure 1D), class 1 antibody 8H12 paired with class 3 antibody 1C4 (Figure 1E), and class 3 antibody 1C4 paired with class 4 antibody 3E2 (Figure 1F).^56^ The dual class 3/class 4 complex (Figure 1F) is especially notable as it reveals how cooperative engagement of distinct epitopes can be achieved through dual targeting of conserved RBD sites.

Class 3 antibodies are characterized by variable lateral face engagement of the RBD, targeting the α2-helix and the β4-β5 hairpin in a manner that does not directly overlap with the ACE2-binding interface. This epitope is accessible in both the “up” and “down” conformations of the RBD and is characterized by moderate conservation across variants. The structural comparison of class 3 antibodies reveals a spectrum of binding modes ranging from most extensive to most compact, with the engagement of the α2-helix serving as the primary variable.

COV2-3835 represents the most extensively engaging class 3 antibody with a footprint spanning the α2-helix (residues 342-349), β4-β5 hairpin (437-449), and RBM loop (463-496).^57^ The antibody engages the RBD through an extensive network of contacts involving all six CDRs, with the heavy chain CDR3 playing a particularly important role in anchoring the antibody to the lateral face. The α2-helix engagement is mediated by critical contacts involving ARG343, THR342, and TYR348, creating a stable interface that effectively immobilizes the lateral face. This extensive engagement translates into greater mechanical constraint on the RBM loop. COV2-3891 exhibits a more compact footprint, spanning residues 437-446 (β4-β5 hairpin) and 498-506 (RBM loop), lacking α2-helix contacts.^57^ This more compact epitope footprint correlates with reduced conformational constraint. In the dual complexes, 1C4 targets a conserved epitope on the lateral face (residues 339, 340, 343-346, 439-448, 450, 451, 499, 500, 509).^117^ By targeting this epitope, 1C4 and 3E2 form an almost contiguous joined footprint covering conserved RBD regions. Importantly, while class 3 antibodies are often described as mediating neutralization through steric hindrance of ACE2 binding^117^, our analysis focuses on the RBD-level interactions and their mechanical consequences, providing a molecular basis for understanding how lateral face engagement can impose constraint on the RBM loop.

In striking contrast to the structural diversity observed among class 3 antibodies, class 4 antibodies target the cryptic inner face of the RBD through a remarkably conserved epitope architecture (Figure 1C-F). The cryptic inner face becomes accessible only when the RBD adopts the “up” conformation, and this epitope is characterized by exceptional conservation across sarbecoviruses, providing a stable anchor for antibody binding that is resistant to mutational escape. The structural analysis of class 4 antibodies 3E2 and COV2-3906 reveals a shared epitope architecture with only subtle variations in footprint. All class 4 antibodies examined in this study share common engagement of three key structural elements that collectively establish the mechanical conduit for long-range allosteric communication. First, the hydrophobic core (residues 364-388) is universally engaged through extensive contacts involving both heavy and light chains. Key residues such as TYR369, PHE377, and THR385 form a stable anchor that explains the extreme rigidification observed at the epitope in the dynamics analysis. The hydrophobic core is structurally rigid and resistant to mutation, providing an anchor for ultra-broad antibody binding while simultaneously serving as the mechanical origin of allosteric signal propagation. Second, the β-sheet core (residues 404-408, 437, 527-534) serves as a conduit for allosteric communication. Residues such as GLY404, ASP405, ARG408, and ASN437 are engaged, allowing conformational changes at the inner face to propagate to the RBM loop through the β-sheet architecture. This engagement establishes the physical pathway through which mechanical changes at the distal inner face are transmitted to the receptor-binding surface. Third, the 370s loop (residues 370-380) is engaged by all class 4 antibodies through main-chain hydrogen bonds that are resistant to mutation, contributing to the exceptional conservation of the class 4 epitope and ensuring stable anchoring of the antibody to the RBD core. While class 4 antibodies share a common epitope architecture, they exhibit subtle variations in their epitope footprints that may influence the magnitude of their allosteric effects. COV2-3906 shows the most extensive engagement of the hydrophobic core and β-sheet core, with all six CDRs making extensive contacts spanning residues 364-388, 527-534, and 434-498, encompassing the hydrophobic core, β-sheet core, and proximal loops. This extensive engagement creates a broad mechanical anchor that can potentially enhance allosteric couplings. 3E2 antibody shows a more focused epitope centered on the hydrophobic core (residues 372-376), β-sheet core (residues 404-408, 437), and proximal loops (498-508). The heavy chain CDR3 inserts into the hydrophobic core, while the light chain contacts the β-sheet core, creating a remarkably stable interface that effectively immobilizes the inner face. In the dual class 3/class 4 complex, the 3E2 epitope is more extensive, with the antibody also engaging the α2-helix (residues 339-346) and β4-β5 hairpin (residues 440-451), suggesting context-dependent expansion of the footprint.

The structural analysis thus reveals that class 3 antibodies exhibit substantial structural diversity in lateral face engagement, with the α2-helix serving as the primary variable determining the extent of mechanical constraint on the RBM loop. In contrast, class 4 antibodies display remarkable structural conservation in their engagement of the cryptic inner face, with the β-sheet core serving as a mechanical conduit for long-range allosteric communication. These distinct epitope architectures provide the foundation for understanding how these antibody classes differentially modulate RBD dynamics.

### Conformational Dynamics of Class 3 and Class 4 Antibody Complexes

To investigate how the distinct structural architectures translate into differential modulation of RBD dynamics, we employed an integrated computational strategy combining CG-CABS simulations with all-atom MD simulations across the six antibody-RBD complexes. This dual-simulation framework enables efficient exploration of conformational space while capturing atomistic detail and solvation effects, ensuring robust characterization of the dynamic response to antibody engagement. From these simulations, we computed residue-specific root-mean-square fluctuations (RMSF), which report directly on the amplitude of thermal motion at each position and reveal how antibody binding locally modulates conformational flexibility across the RBD (Figure 2).

**Figure 2.**
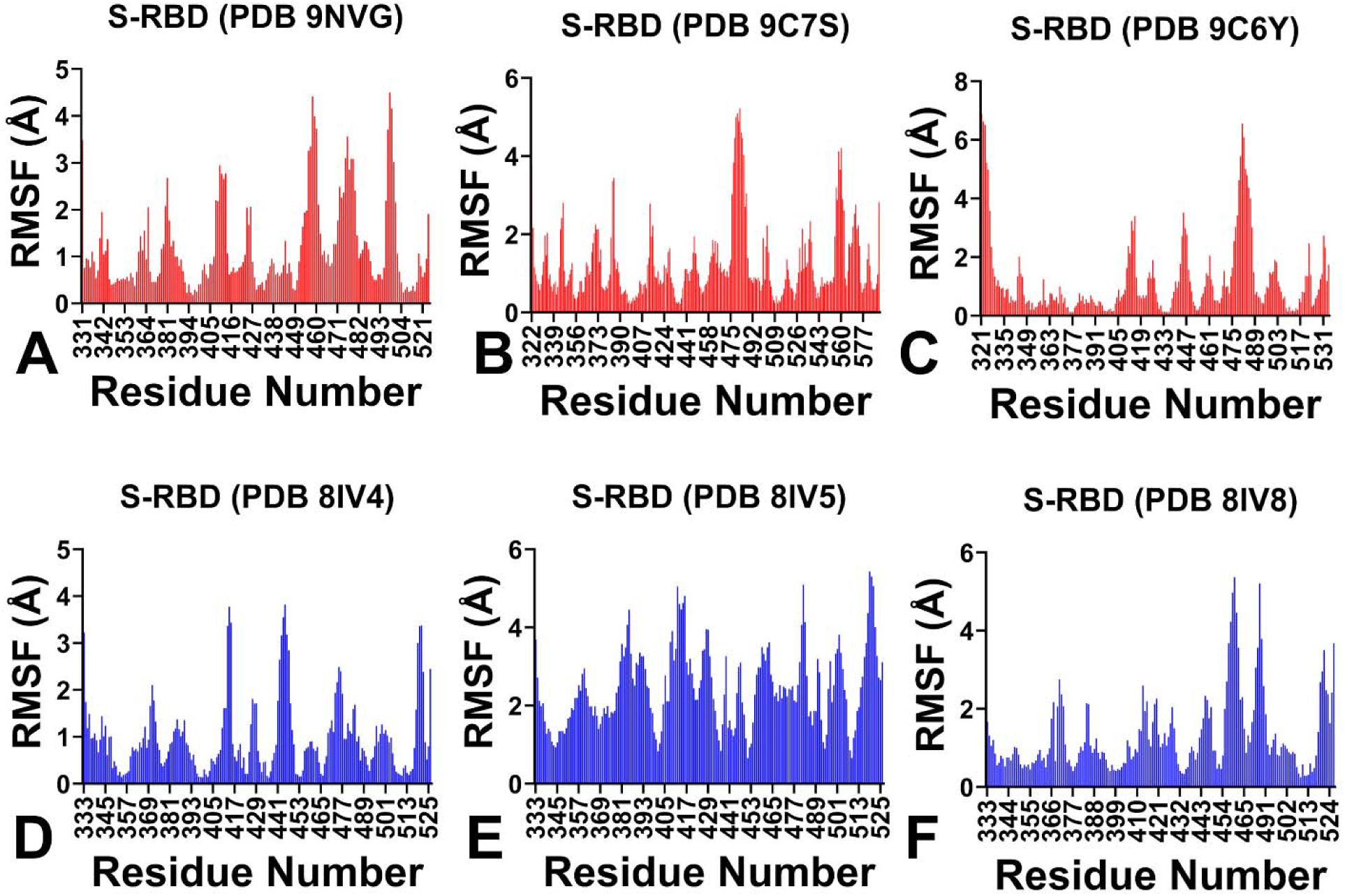
Conformational dynamics of class 3 and class 4 antibody-RBD complexes. RMSF profiles calculated from CG-CABS and all-atom MD simulations across the six antibody-RBD complexes. (A) COV2-3835 (9NVG). (B) COV2-3891 (9C7S). (C) COV2-3906 (9C6Y). (D) 8IV4 complex (class 1 antibody 8H12 + class 4 antibody 3E2). (E) 8IV5 complex (class 1 antibody 8H12 + class 3 antibody 1C4). (F) 8IV8 complex (class 3 antibody 1C4 + class 4 antibody 3E2). The profiles establish the distinct dynamic signatures that distinguish class 3 and class 4 antibody neutralization mechanisms.

The RMSF profiles of the single class 3 antibody complexes (Figure 2A-B) reveal a consistent dynamic signature characterized by pronounced rigidification at the binding interface coupled with moderate constraint of the RBM loop, while regions distal to the epitope retain near-native flexibility. This signature reflects the direct engagement of the lateral face and the antibody’s proximity to the receptor-binding surface. The most rigidified regions correspond precisely to the epitope footprints identified in our structural analysis, demonstrating that the dynamic perturbation is directly localized to the antibody-binding interface. For COV2-3835 (Figure 2A), the RMSF profile reveals relatively low fluctuation values across its broader epitope spanning the α2-helix (residues 342-349), β4-β5 hairpin (437-449), and RBM loop (463-496), with residues such as ARG343, THR342, and TYR348 showing RMSF values below 0.5 Å. This extensive rigidification reflects the comprehensive network of contacts formed by COV2-3835 through all six CDRs. In contrast, COV2-3891 (Figure 2B) exhibits more localized rigidification restricted to its compact β4-β5 hairpin footprint (residues 437-446), with the α2-helix showing significantly higher fluctuation values (RMSF 1.5-2.5 Å). The RBM loop exhibits moderate flexibility, with RMSF values typically ranging from 0.5 to 4.5 Å depending on the specific antibody. The RMSF profiles reveal a spectrum of RBM constraint across the class 3 antibody family that correlates with epitope footprint extent. COV2-3835, with its extensive engagement of the α2-helix and RBM loop, shows the greatest RBM constraint (RMSF values of 2.5-4.5 Å in the RBM loop, residues 470-490), while COV2-3891, with its more compact footprint, exhibits weaker constraint (RMSF values of 2.5-5.5 Å). Notably, regions distal to the binding interface retain significant conformational freedom, suggesting that the dynamic perturbation induced by class 3 antibodies is relatively localized. The class 1/class 3 dual complex (Figure 2E) reveals a dynamic signature characteristic of class 3 engagement, dominated by 1C4’s extensive lateral face contacts with the RBM loop. The lateral face epitope—centered on the β4-β5 hairpin (437-449) and RBM loop (470-490)—shows pronounced rigidification (RMSF <1.0 Å), with the RBM loop showing enhanced constraint (RMSF 1.5-3.0 Å). Residues directly contacted by 1C4 (PHE486, TYR489, GLN493, TYR505) show RMSF values of 1.0-2.0 Å. This signature reflects the class 3 mechanism of localized mechanical constraint without long-range allosteric communication, with the β-sheet core showing minimal perturbation (RMSF 0.5-1.0 Å).

The RMSF profiles of the single class 4 antibody complex (Figure 2C) and the class 1/class 4 dual complex (Figure 2D) reveal a fundamentally different dynamic signature characterized by a stark dichotomy in flexibility patterns: extreme rigidification at the cryptic epitope coupled with pronounced allosteric loosening of the RBM loop. This signature reflects the unique mechanism of class 4 antibodies, which anchor to a structurally rigid hydrophobic core while inducing long-range conformational changes that propagate through the β-sheet core to the RBM. The cryptic epitope—centered on the hydrophobic core (residues 369-385) and the β-sheet core (residues 404-408, 437)—shows remarkably low RMSF values, often below 0.3 Å for key contact residues.

This extreme rigidification is the most pronounced observed across all complexes and reflects the deep, conserved nature of the class 4 epitope, where the antibody forms an extensive network of contacts—including hydrogen bonds with main-chain carbonyl and amide groups—that effectively immobilize the core of the RBD. For COV2-3906 (Figure 2C), the RMSF profile reveals the most extensive rigidification, with hydrophobic core residues TYR369, PHE377, and THR385 showing RMSF values of 0.1-0.2 Å, while the RBM loop exhibits exceptionally high RMSF values of 4.5-5.5 Å. This dramatic gradient from the rigid core to the flexible RBM loop provides direct evidence for long-range allosteric communication through the β-sheet core. The class 1/class 4 dual complex (Figure 2D) reveals a dynamic signature dominated by 3E2’s focused engagement of the hydrophobic core, coupled with localized ACE2-competitive constraint imposed by 8H12 on the RBM loop. The hydrophobic core shows pronounced rigidification (RMSF 0.3-0.6 Å), though less extreme than COV2-3906, reflecting 3E2’s more focused epitope footprint. The RBM loop exhibits enhanced flexibility (RMSF 2.5-4.5 Å), reflecting allosteric loosening induced by 3E2 through the β-sheet core, while the class 1 antibody imposes additional localized constraint (RMSF 2.0-3.5 Å), creating a biphasic dynamic signature. The α2-helix retains significant flexibility (RMSF 1.5-2.5 Å), consistent with the absence of class 3 antibody contacts.

The most intriguing system is the dual class 3/class 4 complex (Figure 2F), which shows a dramatically amplified dynamic signature resulting from simultaneous engagement of both antibody classes on non-overlapping epitopes. The hydrophobic core shows the most extreme rigidification observed across all complexes (RMSF <0.2 Å for TYR369, PHE377, THR385), reflecting the combined anchoring effect of 3E2 and the broader structural stabilization from 1C4. The β-sheet core residues GLY404, ASP405, ARG408, and ASN437 show RMSF values of 0.2-0.4 Å, indicating enhanced mechanical coupling through the allosteric conduit. Paradoxically, despite direct engagement of the RBM loop by 1C4, the RBM loop exhibits exceptionally high RMSF values exceeding 5.0 Å for key residues such as TYR489, GLN493, and TYR505. This observation—that direct engagement of the RBM loop does not result in constraint but rather enhanced flexibility—reflects the amplified allosteric communication established by dual engagement. The combination of 3E2 anchoring to the hydrophobic core and 1C4 engagement of the β4-β5 hairpin creates a more extensive footprint that enhances propagation of conformational changes through the β-sheet core, resulting in amplified RBM loosening that overcomes the direct steric constraint imposed by 1C4.

To complement the RMSF analysis, we calculated the Distance Fluctuation Stability Index (DFSI) for each residue across the six antibody-RBD complexes (Figure 3). The DFSI, defined as the inverse of the mean-square fluctuation of each residue, provides a direct measure of the effective force constant governing local positional deviations. Higher DFSI values indicate greater mechanical rigidity and stronger local restoring forces, while lower values reflect increased flexibility. The DFSI profiles offer a distinct and complementary perspective from the RMSF analysis: while RMSF identifies regions that become rigidified or mobilized upon antibody binding, DFSI quantifies the magnitude of these mechanical changes in terms of local stiffness, revealing the depth of mechanical perturbation and the gradient of rigidity propagation through the protein structure. The DFSI profiles for the single class 3 antibody complexes (Figure 3A-B) reveal that mechanical stiffening is strictly confined to the antibody-binding interface, with the highest values concentrated at residues directly contacted by the antibody. For COV2-3835 (Figure 3A), the elevated DFSI values extend across the entire epitope footprint— encompassing the α2-helix (residues 342-349), β4-β5 hairpin (437-449), and RBM loop (463-496)—with the most pronounced stiffening observed at residues ARG343, THR342, and TYR348. The RBM loop shows intermediate DFSI values, reflecting the moderate mechanical constraint imposed by direct antibody contacts. However, regions distal to the binding interface exhibit the lowest DFSI values, comparable to those of the unbound RBD, confirming that the mechanical perturbation does not propagate beyond the immediate epitope.

**Figure 3.**
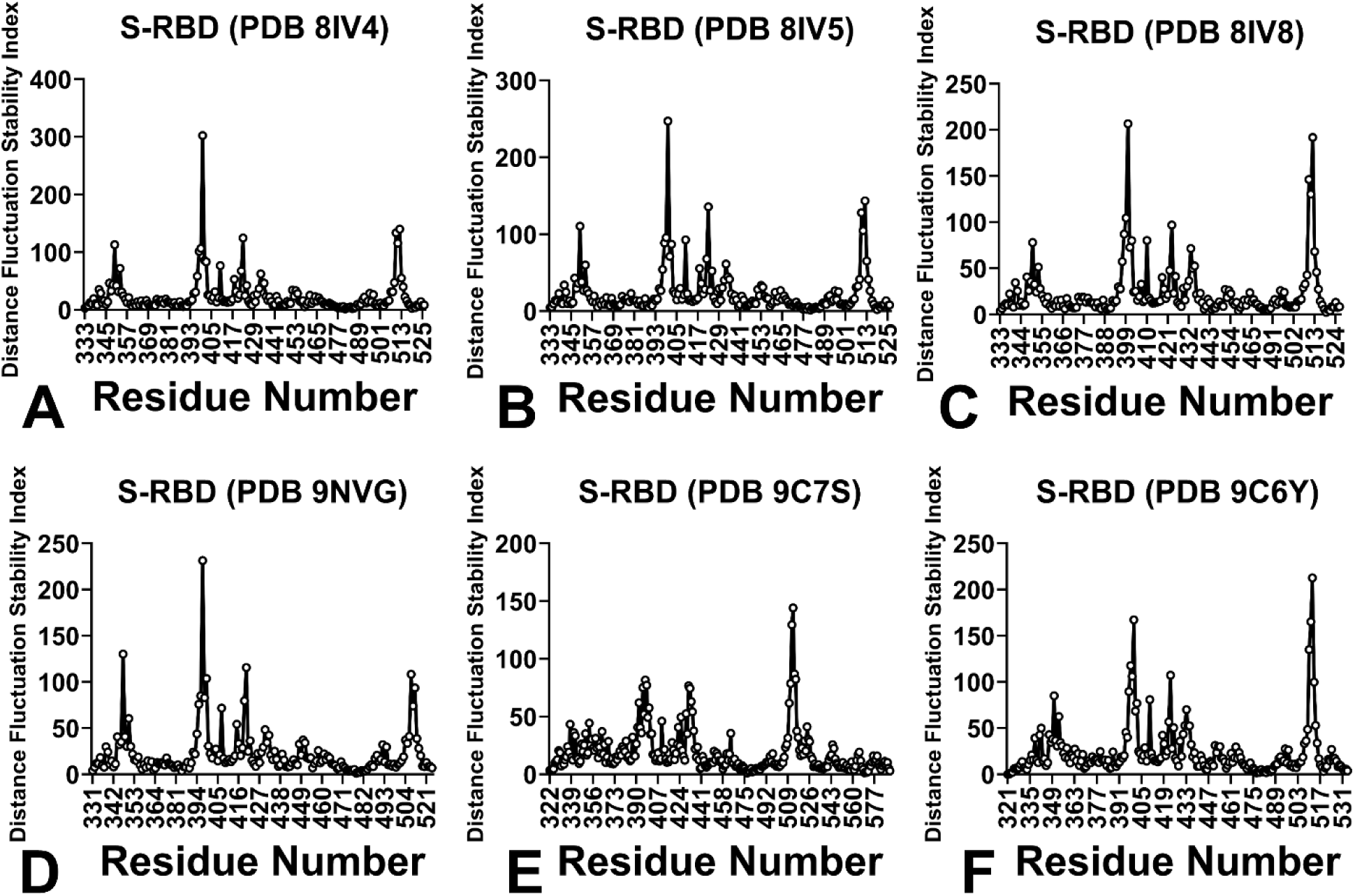
Distance Fluctuation Stability Index (DFSI) analysis of antibody-RBD complexes. Quantitative mechanical rigidity profiling across the six antibody-RBD complexes. Higher DFSI values indicate greater mechanical rigidity and stronger local restoring forces. (A) COV2-3835 (9NVG). (B) COV2-3891 (9C7S). (C) COV2-3906 (9C6Y). (D) 8IV4 complex (class 1 antibody 8H12 + class 4 antibody 3E2). (E) 8IV5 complex (class 1 antibody 8H12 + class 3 antibody 1C4). (F) 8IV8 complex (class 3 antibody 1C4 + class 4 antibody 3E2). The DFSI profiles provide a quantitative mechanical map distinguishing localized stiffening (class 3) from long-range mechanical communication (class 4).

In contrast, COV2-3891 (Figure 3B) exhibits a markedly different DFSI signature, with stiffening restricted to the compact β4-β5 hairpin footprint (residues 437-446), while the α2-helix shows minimal mechanical perturbation. The RBM loop shows reduced DFSI values compared to COV2-3835, reflecting the more limited conformational constraint. The class 1/class 3 dual complex (Figure 3E) reveals enhanced mechanical stiffening in the RBM loop compared to the class 1/class 4 complex (Figure 3D), reflecting the direct steric constraint imposed by 1C4.

The DFSI profiles for the single class 4 antibody complex (Figure 3C) and the class 1/class 4 dual complex (Figure 3D) reveal a dramatic mechanical dichotomy that is the defining signature of allosteric communication. The hydrophobic core exhibits exceptionally high DFSI values, indicating extreme mechanical rigidity that reflects the deep, conserved nature of the class 4 epitope. This extreme stiffening is most pronounced at residues TYR369, PHE377, and THR385. For COV2-3906 (Figure 3C), the hydrophobic core shows the highest DFSI values among all class 4 complexes, consistent with its most extensive engagement of the hydrophobic core and β-sheet core through all six CDRs. The β-sheet core also shows substantially elevated DFSI values, consistent with its role as the mechanical conduit for allosteric signal propagation, with residues GLY404, ASP405, ARG408, and ASN437 showing pronounced stiffening. Critically, these elevated DFSI values in the core are accompanied by dramatically reduced DFSI values in the RBM loop, representing a marked reduction in mechanical rigidity compared to the core epitope. This stark contrast between the rigid core and the flexible RBM loop provides direct quantitative evidence for long-range allosteric communication, demonstrating that the engagement of the β-sheet core by class 4 antibodies allows mechanical changes at the inner face to propagate to the receptor-binding surface. The DFSI profile reveals a clear gradient from the extremely rigid core to the dramatically loosened RBM loop, with the β-sheet core serving as the mechanical conduit.

The class 1/class 4 dual complex (Figure 3D) shows a more moderate DFSI signature, with the hydrophobic core exhibiting elevated DFSI values but reduced magnitude compared to COV2-3906, reflecting 3E2’s focused engagement. The β-sheet core shows intermediate DFSI values, indicating less extensive mechanical coupling, while the RBM loop displays DFSI values that reflect the balance between allosteric loosening from 3E2 and direct constraint from 8H12. This intermediate mechanical signature demonstrates that the magnitude of allosteric communication is governed by the extent of inner-face engagement, with more extensive footprints yielding stronger mechanical gradients.

The dual class 3/class 4 complex (Figure 3F) reveals the most extreme mechanical gradient observed across all complexes. The hydrophobic core exhibits the highest DFSI values, reflecting the combined anchoring effect of 3E2 and the broader structural stabilization from 1C4. The β-sheet core shows enhanced DFSI values compared to individual antibody complexes, indicating amplified mechanical coupling through the allosteric conduit. Remarkably, the RBM loop exhibits the lowest DFSI values observed across all complexes, despite direct engagement by 1C4. This observation that direct engagement of the RBM loop does not result in mechanical stiffening but rather amplified loosening reflects the dominance of the allosteric communication pathway established by cooperative inner-face and lateral-face engagement. The dual engagement creates a more extensive footprint that enhances the propagation of mechanical changes through the β-sheet core, resulting in amplified RBM mechanical loosening that overcomes the direct steric constraint.

The integration of RMSF and DFSI profiles provides a comprehensive and physically interconnected view of how class 3 and class 4 antibodies differentially modulate RBD mechanics. RMSF profiles identify which regions become rigidified or mobilized, while DFSI profiles quantify the magnitude of these mechanical changes in terms of local stiffness, revealing how antibody-induced perturbations propagate through the protein structure. Together, these complementary measures establish that class 3 antibodies induce a localized mechanical perturbation where stiffening is confined to the binding interface, with limited propagation to distal regions. The correlation between epitope footprint extent and the magnitude of mechanical stiffening across class 3 antibodies establishes the mechanical state of the RBM loop as a key determinant of neutralization efficacy. Class 4 antibodies establish a long-range mechanical gradient from extreme stiffening at the hydrophobic core to pronounced loosening at the RBM loop, with the β-sheet core serving as the essential mechanical conduit. The magnitude of this mechanical gradient correlates with the extent of inner-face engagement, with more extensive footprints yielding stronger allosteric effects.

### Mutational Profiling of RBD-Antibody Binding Interfaces

To systematically evaluate the molecular determinants of immune sensitivity and escape vulnerability, we constructed mutational heatmaps for the RBD interface residues of all six antibody-RBD complexes (Figures 4, 5). Using the BeAtMuSiC approach for mutational scanning, we assessed how amino acid substitutions at each RBD position impact antibody binding affinity, providing quantitative insights into epitope sensitivity and the potential for viral escape. The heatmaps display the computed binding free energy changes (ΔΔG) for all 20 single amino acid substitutions at each interface residue. The mutational heatmaps for the single class 3 antibody complexes (Figure 4A-B) reveal a spectrum of sensitivity patterns that reflect the structural diversity observed within this antibody class. All class 3 antibodies share a common mutational signature at the β4-β5 hairpin (residues 437-449), which serves as a structural anchor for the lateral face epitope. Residues 446-449 consistently show significant mutational sensitivity (ΔΔG > 1.5 kcal/mol), reflecting their critical role in stabilizing the antibody-RBD interface. This shared sensitivity confirms the β4-β5 hairpin as the core anchor for class 3 antibodies, providing a stable contact point that is conserved across the class. In contrast to the conserved β4-β5 hairpin, the α2-helix (residues 340-360) and RBM loop (residues 470-490) show variable mutational sensitivity that reflects the structural plasticity of class 3 antibody engagement.

**Figure 4.**
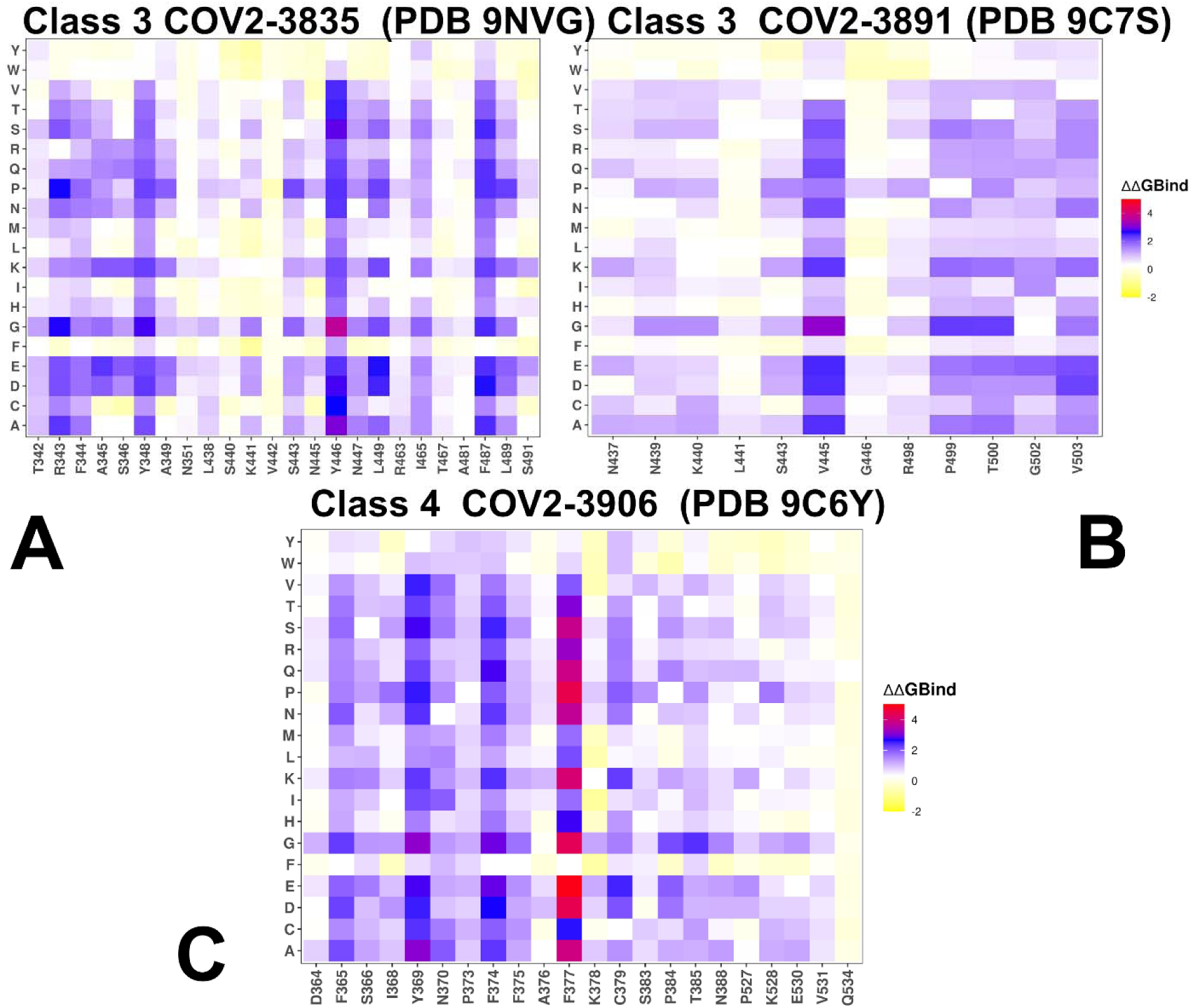
Mutational profiling of RBD-antibody binding interfaces. Mutational heatmaps displaying computed binding free energy changes (ΔΔG) for all 20 single amino acid substitutions at each RBD interface residue across antibody-RBD complexes (A) class 3 COV2-3835 (PDB 9NVG), (B) class 3 COV2-3891 (PDB 9C7S) and (C) class 4 COV2-3906 (PDB 9C6Y). The binding energy hotspots correspond to residues with high mutational sensitivity. The squares on the heatmap are colored using a 4-colored scale yellow-white-blue-red with blue-red indicating the largest unfavorable effect on binding and stability (ΔΔG > 2.0 kcal/mol), while yellow-white points to mutations that have neutral or favorable effect and improve binding. The standard errors of the mean for binding free energy changes using randomly selected 1,000 conformational samples (0.08-0.12 kcal/mol) obtained from the atomistic trajectories.

**Figure 5.**
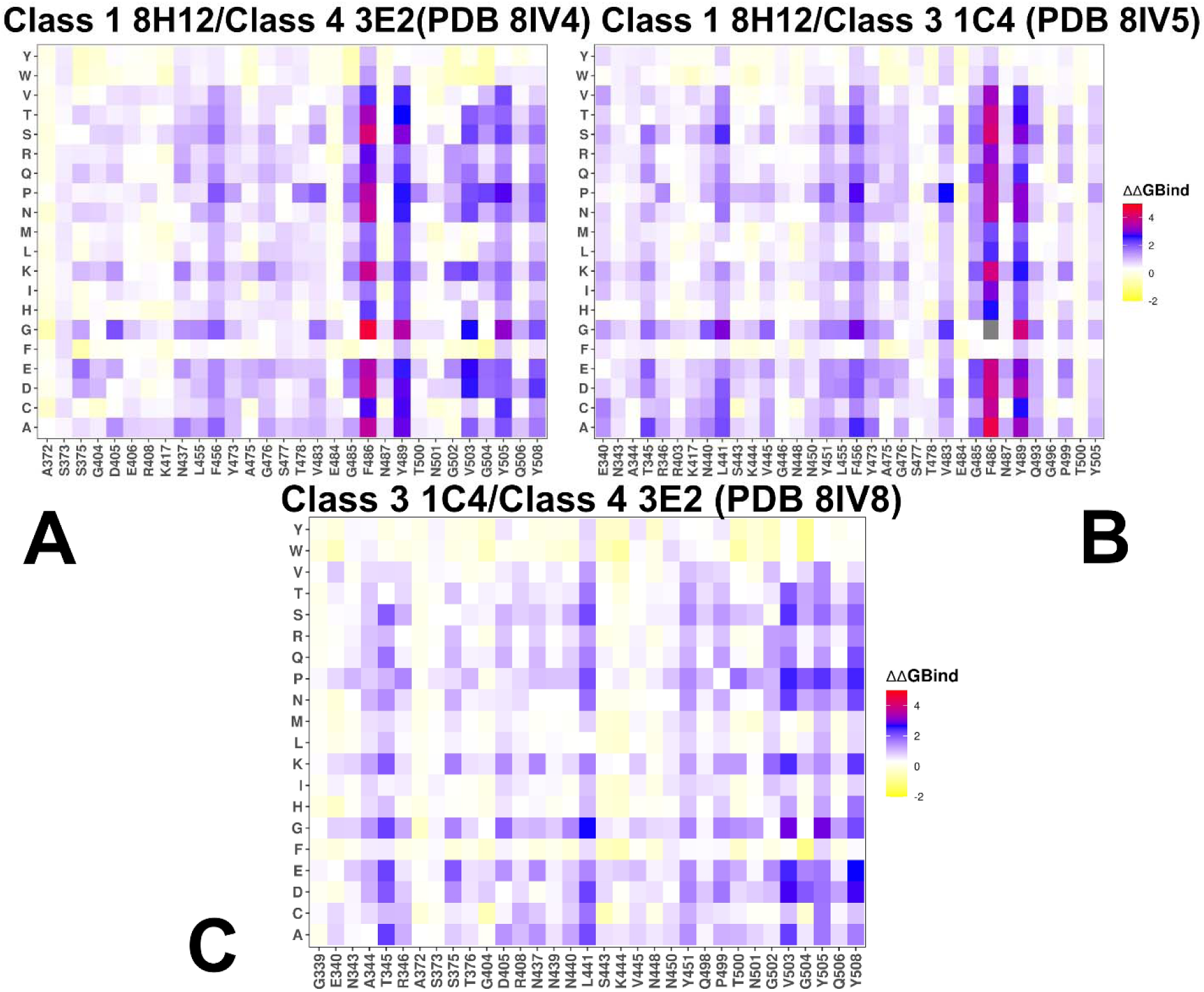
Mutational profiling of RBD-antibody binding interfaces. Mutational heatmaps displaying computed binding free energy changes (ΔΔG) for all 20 single amino acid substitutions at each RBD interface residue across antibody-RBD complexes (A) Dual complex of class 1 antibody 8H12 and class 4 antibody 3E2 (PDB 8IV4). (B) Dual complex of class 1 antibody 8H12 and class 3 antibody 1C4 (PDB 8IV5). (C) Dual complex of class 3 antibody 1C4 and class 4 antibody 3E2 (PDB 8IV8). The binding energy hotspots correspond to residues with high mutational sensitivity. The squares on the heatmap are colored using a 4-colored scale yellow-white-blue-red with blue-red indicating the largest unfavorable effect on binding and stability (ΔΔG > 2.0 kcal/mol), while yellow-white points to mutations that have neutral or favorable effect and improve binding. The standard errors of the mean for binding free energy changes using randomly selected 1,000 conformational samples (0.06-0.15 kcal/mol) obtained from the atomistic trajectories.

For COV2-3835 (Figure 4A), the heatmap reveals the most extensive mutational sensitivity among class 3 antibodies, with critical hotspots spanning the α2-helix (residues 342-349: THR342, ARG343, TYR348), β4-β5 hairpin (437-449), and RBM loop (463-496). Residues ARG343, THR342, TYR348, Y446, and F487 show particularly intense sensitivity (ΔΔG > 2.5 kcal/mol), consistent with their role in anchoring the antibody to the α2-helix. In contrast, COV2-3891 (Figure 4B) shows a more focused pattern centered on the β4-β5 hairpin (residues 437-446) and the proximal RBM loop (residues 498-506). The α2-helix shows minimal sensitivity (ΔΔG < 0.5 kcal/mol), consistent with COV2-3891’s compact footprint. The key hotspots for this antibody are V445, P499, T500, G502, and V503. The differential sensitivity directly correlates with the structural differences observed between these two antibodies, with the more extensive COV2-3835 epitope creating a broader set of sequence constraints.

The class 1/class 3 dual complex (Figure 5B) reveals enhanced mutational sensitivity consistent with the broader epitope footprint of 1C4. The heatmap shows intense sensitivity spanning the RBM loop (455-496; ΔΔG > 2.0 kcal/mol) and β4-β5 hairpin (437-449; ΔΔG > 1.5 kcal/mol). Hotspot residues such as LEU455, PHE456, PHE486, and TYR489 show significant mutational sensitivity (ΔΔG 1.5-2.5 kcal/mol). The β4-β5 hairpin anchor remains consistently sensitive (ΔΔG > 1.5 kcal/mol), confirming its role as the invariant core of class 3 epitope recognition. The enhanced sensitivity at the RBM loop in this dual complex compared to the single antibody complexes reflects the combined effect of the class 3 and class 1 antibodies, which together impose a more extensive set of sequence constraints on the receptor-binding surface.

The mutational heatmaps for the class 4 antibody complexes (Figure 4C, Figure 5A, Figure 5C) reveal a fundamentally different and remarkably consistent pattern characterized by an invariant hydrophobic core that is exquisitely sensitive to mutation. This core, spanning residues 369-385, represents the most mutationally constrained region across all complexes examined, with essentially every position showing intense red/orange coloring. The heatmaps consistently highlight TYR369, PHE377, C379, Y380, and THR385 as the most sensitive positions, with virtually any substitution severely compromising antibody binding (ΔΔG > 3.0 kcal/mol). These residues form the structural heart of the class 4 epitope, providing the hydrophobic surface that anchors the antibody. The invariance of these residues across SARS-CoV-2 variants reflects their essential role for antibody recognition and structural integrity of the RBD fold. Adjacent to the hydrophobic core, the β-sheet core (residues 404-408, 437) forms a second tier of critical contacts. GLY404, ASP405, ARG408, and ASN437 consistently show intense mutational sensitivity (ΔΔG > 2.0 kcal/mol), reflecting their role as the structural conduit for allosteric communication. These residues connect the inner face to the RBM through the β-sheet architecture, and their conservation is essential for maintaining the long-range communication pathway that class 4 antibodies exploit. The 370s loop and proximal loops (498-508) show a more nuanced pattern, with a “gradient of sensitivity” from the absolutely invariant core to the moderately tolerant periphery. For the single class 4 antibody complex (Figure 4C), the heatmap shows the most extensive mutational sensitivity among class 4 antibodies, with intense red/orange coloring spanning the entire hydrophobic core (residues 364-388), β-sheet core, and proximal loops. Key hotspots include TYR369, F374, F375, S383, and T385. The broad sensitivity across the entire epitope explains the exceptional binding affinity and ultra-broad neutralization of COV2-3906.

The class 1/class 4 dual complex (Figure 5A) shows a more moderate mutational sensitivity pattern with pronounced hotspots at conserved positions F486, Y489, V503, Y505, and Y508. The dual class 3/class 4 complex (Figure 5C) reveals the most extensive sensitivity among all complexes, with critical hotspots spanning the hydrophobic core (369-385), β-sheet core (404-408, 437), and RBM loop (455-496). Notably, the RBM loop residues PHE486, TYR489, GLN493, and TYR505 show significant mutational sensitivity (ΔΔG 1.5-2.5 kcal/mol) in this dual complex, indicating that 1C4’s direct engagement of the RBM loop creates additional sequence constraints beyond those imposed by the hydrophobic core alone.

However, the hydrophobic core residues remain the most sensitive (ΔΔG > 3.0 kcal/mol), confirming their role as the immutable anchor of class 4 recognition. The comparison between computational predictions and DMS experimental data reveals excellent agreement, with our mutational scanning accurately recapitulating the known binding hotspots and escape mutations for both antibody classes.^42–45,118,119^ For class 3 antibodies, the foundational hotspot residues forming the core anchor sit on a loop-dense, accessible outer shelf including R346, K444, and N450. DMS profiles identified K417, L452, and V445 as important mutational centers. For class 4 antibodies, the most critical binding residues are grouped tightly along the RBD short cliff and correspond to C379, Y380, V382, and T385. Some class 4 antibodies approach the inner face at a sharper perpendicular angle, producing secondary hotspots on the upper border of the hidden core (V405, G502, V503, G504, and V505 residues).

### MM-GBSA Binding Free Energy Analysis: Energetic Architecture and Hotspot Characterization

To complement the mutational scanning analysis and provide a quantitative, physically grounded characterization of binding affinity hotspots, we performed MM-GBSA binding free energy calculations with residue-based energy decomposition across all six antibody-RBD complexes. This approach partitions the total binding free energy into van der Waals (VDW) and electrostatic (ELE) contributions at the per-residue level (Supporting Figures S1-S2), revealing the physical nature of the interactions that define the binding affinity hotspots identified in our mutational heatmaps. The total binding energy profiles (Figure 6) reveal a clear hierarchy of energetic contributions that distinguishes the two antibody classes and explains their differential sensitivity to mutational perturbation.

**Figure 6.**
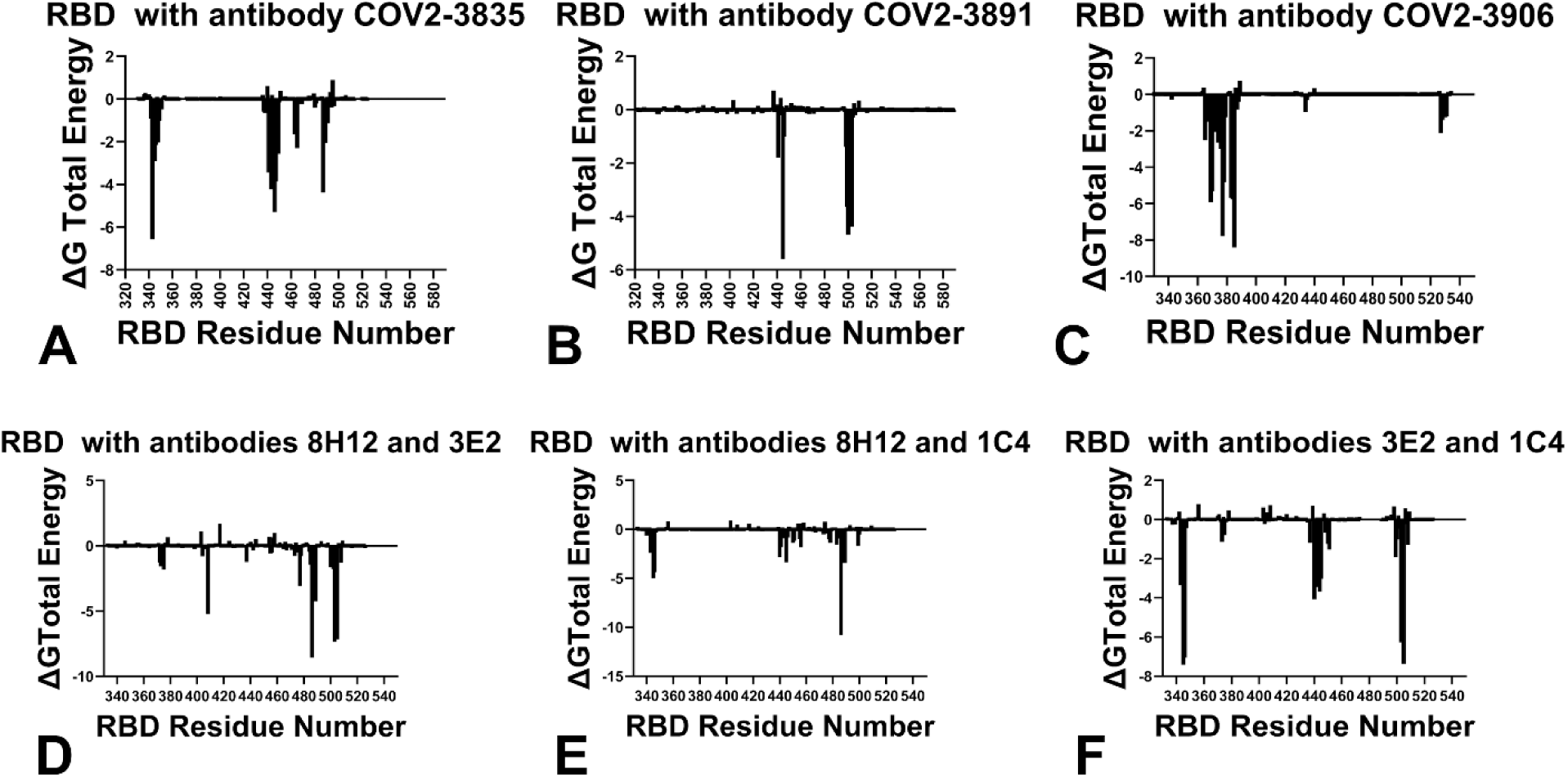
MM-GBSA binding free energy analysis. Total binding energy profiles per RBD residue across the six antibody-RBD complexes, revealing the energetic architecture and binding affinity hotspots underlying the distinct neutralization mechanisms. (A) COV2-3835 (9NVG). (B) COV2-3891 (9C7S). (C) COV2-3906 (9C6Y). (D) 8IV4 complex (class 1 antibody 8H12 + class 4 antibody 3E2). (E) 8IV5 complex (class 1 antibody 8H12 + class 3 antibody 1C4). (F) 8IV8 complex (class 3 antibody 1C4 + class 4 antibody 3E2). The profiles reveal a clear hierarchy of energetic contributions distinguishing class 3 (VDW-driven binding distributed across multiple hotspots) from class 4 (extreme VDW stabilization at the hydrophobic core reinforced by main-chain electrostatic interactions).

The MM-GBSA energy decomposition for the single class 3 antibody complexes reveals that binding is predominantly driven by favorable van der Waals interactions (Supporting Figure S1A-B), with electrostatic contributions (Supporting Figure S2A-B) playing a secondary, often context-dependent role. For COV2-3835 (Figure 6A), the per-residue energy decomposition identifies a set of primary binding hotspots that precisely correspond to the mutational sensitivity peaks observed in the heatmaps. The most favorable total binding energy contributions are concentrated at the α2-helix (residue ARG343; ΔG_total = −6.57 kcal/mol), the β4-β5 hairpin anchor (residues LYS437, VAL442, GLY446; ΔG_total = −0.59 to −1.17 kcal/mol), and the RBM loop (residue PHE487; ΔG_total = −4.38 kcal/mol). The VDW contributions (Supporting Figure S1A) are the dominant driver of binding, with energies of −4.0 to −6.0 kcal/mol at the α2-helix and RBM loop, and −0.5 to −2.0 kcal/mol at the β4-β5 hairpin. The electrostatic contributions (Supporting Figure S2A) are more variable, with ARG343 showing favorable interactions (−46.70 kcal/mol) that are partially offset by solvation (49.74 kcal/mol), resulting in a net total contribution of −6.57 kcal/mol. Other residues such as LYS441 show favorable electrostatic interactions (−50.97 kcal/mol) with partial solvation compensation. Notably, residue ASN437 (referred to as ASN434 in the 9NVG structure) shows a total binding energy contribution of approximately −1.25 kcal/mol, consistent with its role in stabilizing the β4-β5 hairpin anchor. For COV2-3891 (Figure 6B), the energy decomposition reveals a more focused pattern consistent with its compact epitope footprint. The β4-β5 hairpin residues (VAL445; ΔG_total = −5.59 kcal/mol; THR444; ΔG_total = −4.69 kcal/mol) serve as the primary energetic hotspots, while the α2-helix shows negligible total binding energy consistent with the absence of direct antibody contacts in this region. The RBM loop residues (498-506) show moderate contributions (ΔG_total = −3.0 to −4.0 kcal/mol), reflecting the more limited engagement compared to COV2-3835. The VDW contributions (Supporting Figure S1B) confirm the β4-β5 hairpin as the primary anchor (−4.0 to −6.0 kcal/mol), with the α2-helix showing minimal VDW contributions (−0.5 to −1.0 kcal/mol). Electrostatic contributions are minimal across the epitope (Supporting Figure S2B), with no residues showing favorable interactions exceeding −1.0 kcal/mol. This differential energetic architecture—with COV2-3835 distributing binding energy across multiple regions while COV2-3891 concentrates energy at a single anchor—explains the observed differences in mutational sensitivity and neutralization potency between these two class 3 antibodies.

The class 1/class 3 dual complex (Figure 6E) reveals context-dependent modulation of the energetic landscape. The total binding energy profile shows substantial stabilization at the RBM loop residues (PHE486; ΔG_total = −10.83 kcal/mol; TYR489; ΔG_total = −3.44 kcal/mol) and the β4-β5 hairpin (VAL445; ΔG_total = −3.37 kcal/mol; GLY446; ΔG_total = −0.85 kcal/mol). The VDW contributions (Supporting Figure S1E) show enhanced stabilization at the RBM loop (−9.45 kcal/mol for PHE486) and β4-β5 hairpin (−2.83 kcal/mol for VAL445), while electrostatic contributions (Supporting Figure S2E) show variable contributions that are partially compensated by solvation. This energetic architecture explains the enhanced neutralization potency of the 1C4-8H12 antibody pair, which achieves high affinity through distributed stabilization across the RBM loop and β4-β5 hairpin.

The MM-GBSA energy decomposition for the class 4 antibody complexes reveals a fundamentally different energetic architecture, characterized by extreme VDW stabilization at the hydrophobic core coupled with extensive main-chain electrostatic interactions that confer exceptional resistance to mutational escape. For the single class 4 antibody (Figure 6C), the per-residue decomposition identifies the hydrophobic core residues as the dominant energetic hotspots. The total binding energy profile reveals that the hydrophobic core residues (TYR369; ΔG_total = −5.92 kcal/mol; PHE377; ΔG_total = −7.79 kcal/mol; THR385; ΔG_total = −8.41 kcal/mol) contribute the most favorable energies, while the β-sheet core provides secondary stabilization (PRO384; ΔG_total = −5.77 kcal/mol; ASN370; ΔG_total = −5.33 kcal/mol; SER383; ΔG_total = −5.70 kcal/mol). The VDW contributions (Supporting Figure S1C) reveal extreme stabilization at the hydrophobic core (−7.37 kcal/mol for TYR369; −7.59 kcal/mol for PHE377; −2.37 kcal/mol for THR385), reflecting the dense hydrophobic packing that anchors the antibody to the structurally rigid core and explaining why mutations at these positions are catastrophic for binding (ΔΔG > 3.0 kcal/mol in mutational heatmaps). The β-sheet core residues (PRO384; ΔG_total = −5.77 kcal/mol; ASN370; ΔG_total = −5.33 kcal/mol; SER383; ΔG_total = −5.70 kcal/mol) show substantial contributions, consistent with their role as structural conduits for allosteric communication. Notably, the electrostatic contributions (Supporting Figure S2C) reveal extensive main-chain hydrogen bonds involving the 370s loop and β-sheet core, with residues N370, K378, S383, and T385 showing favorable electrostatic interactions (−17.25 kcal/mol for N370; −65.59 kcal/mol for K378; −11.95 kcal/mol for S383; −15.52 kcal/mol for THR385) that are partially offset by solvation. These main-chain electrostatic interactions are resistant to mutation, as they depend on the backbone conformation rather than side-chain identity, contributing to the exceptional conservation of the class 4 epitope. The total binding energy for COV2-3906 is −133.77 kcal/mol, the most favorable among single-antibody complexes, consistent with its ultra-broad neutralization profile.

The class 1/class 4 dual complex (Figure 6D) shows a more moderate energetic signature. The hydrophobic core residues (ALA372; ΔG_total = −1.27 kcal/mol; SER373; ΔG_total = −1.60 kcal/mol; SER375; ΔG_total = −1.83 kcal/mol) serve as the primary hotspots, while the β-sheet core (ARG408; ΔG_total = −5.24 kcal/mol) provides secondary stabilization. The RBM loop residues (PHE486; ΔG_total = −8.57 kcal/mol; TYR489; ΔG_total = −4.24 kcal/mol) show substantial contributions, reflecting the additional engagement of the class 1 antibody 8H12. The VDW contributions (Supporting Figure S1D) show −9.27 kcal/mol at PHE486 and −5.05 kcal/mol at TYR489, while electrostatic contributions (Supporting Figure S2D) show variable contributions with significant solvation compensation. The total binding energy for the class 1/class 4 dual complex is −89.70 kcal/mol, reflecting the combined stabilization from both antibodies.

The dual class 3/class 4 complex (Figure 6F) reveals the most extensive energetic stabilization among all complexes. The total binding energy profile shows extreme stabilization at the hydrophobic core (THR345; ΔG_total = −7.42 kcal/mol; ARG346; ΔG_total = −7.06 kcal/mol; TYR505; ΔG_total = −7.38 kcal/mol), enhanced stabilization at the β-sheet core (LYS444; ΔG_total = −3.68 kcal/mol; VAL445; ΔG_total = −3.02 kcal/mol; ASN343; ΔG_total = −3.36 kcal/mol), and significant contributions at the RBM loop (LEU441; ΔG_total = −3.45 kcal/mol; TYR505; ΔG_total = −7.38 kcal/mol). The VDW contributions (Supporting Figure S1F) reveal - 4.26 kcal/mol at THR345, −3.70 kcal/mol at ARG346, and −4.00 kcal/mol at TYR505, reflecting the combined anchoring effect of 3E2 and the broader structural stabilization from 1C4. The electrostatic contributions (Supporting Figure S2F) show extensive main-chain hydrogen bonds involving the 370s loop and β-sheet core (ARG346; −40.03 kcal/mol; LYS444; −40.84 kcal/mol), with significant solvation compensation. The total binding energy for the dual class 3/class 4 complex is −106.96 kcal/mol, reflecting the most extensive energetic stabilization among all complexes, yet the neutralization potency remains limited by the inherent inefficiency of the allosteric mechanism.

The integration of MM-GBSA energy decomposition with the mutational heatmaps provides a powerful validation of our hotspot predictions and reveals the physical basis for the observed mutational sensitivity patterns. Across all complexes, there is excellent agreement between the residues showing the largest total binding energy contributions (Figure 6) and those identified as primary binding hotspots in the mutational scanning (Figures 4,5). For class 3 antibodies, the residues with the largest total energy contributions (ARG343; −6.57 kcal/mol, and PHE487; −4.38 kcal/mol in the single class 3 COV2-3835 complex; VAL445; −5.59 kcal/mol, and THR444; −4.69 kcal/mol in the single class 3 COV2-3891 complex; PHE486; −10.83 kcal/mol, and TYR489; −3.44 kcal/mol in the class 1/class 3 dual complex) are precisely those showing the highest mutational sensitivity (ΔΔG > 2.0 kcal/mol). This agreement confirms that the mutational sensitivity reflects the underlying energetic importance of these residues, with VDW-driven hydrophobic packing serving as the primary determinant of binding affinity. For class 4 antibodies, the residues with the largest total energy contributions (THR385; −8.41 kcal/mol, PHE377; −7.79 kcal/mol, and TYR369; −5.92 kcal/mol in the single class 4 complex; PHE486; - 8.57 kcal/mol, and ARG408; −5.24 kcal/mol in the class 1/class 4 dual complex; TYR505; −7.38 kcal/mol, THR345; −7.42 kcal/mol, and ARG346; -7.06 kcal/mol in the dual class 3/class 4 complex) correspond to the residues showing the most extreme mutational sensitivity (ΔΔG > 3.0 kcal/mol). The electrostatic contributions further explain why certain residues show even greater sensitivity than predicted by VDW alone—the main-chain hydrogen bonds involving N370, K378, S383, and T385 create a network of interactions that is both energetically important and resistant to mutation, as mutations that disrupt these interactions would require backbone conformational changes that are structurally disfavored. This explains the exceptional conservation of the class 4 epitope and the limited escape pathways available to the virus.

The comparison between computational predictions and experimental DMS data further validates our approach. For class 3 antibodies, the predicted hotspots at the β4-β5 hairpin (LYS437, VAL445, GLY446) and α2-helix (ARG343, THR342) correspond to experimentally identified escape residues where mutations are observed in circulating variants. The predicted hotspots at the RBM loop (PHE486, TYR489) correspond to residues where mutations are known to affect ACE2 binding, explaining why these positions are under dual selective pressure—mutations that escape antibody binding may also impair receptor binding, limiting their fitness. For class 4 antibodies, the predicted hotspots at the hydrophobic core (TYR369, PHE377, THR385) and β-sheet core (GLY404, ASP405, ARG408) correspond to positions that are highly conserved and where escape mutations are rarely observed in circulating variants due to prohibitive fitness costs. The main-chain electrostatic interactions identified in our MM-GBSA analysis explain why these residues are so resistant to mutation—they are essential for maintaining the β-sheet architecture that is critical for RBD structural integrity.

The MM-GBSA energy decomposition provides a quantitative framework for predicting antibody resistance and guiding therapeutic design. For class 3 antibodies, the energetic hierarchy suggests that engineering strategies should focus on enhancing interactions at the β4-β5 hairpin anchor while optimizing contacts at the α2-helix and RBM loop to achieve both high potency and resilience. Antibodies that distribute binding energy across multiple hotspots are likely to be more resilient to escape than those that concentrate energy at a single anchor as mutations that disrupt one hotspot can be compensated by persistent contacts at others. For class 4 antibodies, the energetic hierarchy suggests that the hydrophobic core is effectively immutable, providing a stable anchor that ensures ultra-broad binding. However, the allosteric mechanism limits neutralization potency, suggesting that strategies to enhance the allosteric effect—such as engineering multivalent antibodies that engage multiple regions of the inner face (dual class 3/class 4 complex-like)—could improve efficacy while maintaining the high genetic barrier to resistance. The main-chain electrostatic interactions at the 370s loop and β-sheet core further suggest that antibodies targeting these regions may be particularly resistant to escape, as mutations that disrupt these interactions would require backbone conformational changes that are structurally disfavored.

### Ensemble-Based Allosteric Network Analysis of the RBD-Antibody Complexes

To elucidate the allosteric communication networks that underlie the distinct neutralization mechanisms, we performed residue interaction network analysis across all six antibody-RBD complexes (Figure 7). This graph-based approach models the protein as a network of residues connected by non-covalent interactions, enabling the identification of key communication hubs that mediate long-range signal propagation. We computed two complementary network parameters that together reveal the architecture of allosteric communication pathways. The Short Path Betweenness (SPC) quantifies the extent to which a residue serves as a critical conduit for communication between distant regions of the protein—high SPC values indicate that a residue lies on many shortest paths connecting other residue pairs, establishing it as a structural hub within the communication network. The Z-score of Average Short Path Length (Z-ASPL) identifies residues whose mutational perturbation significantly alters the global communication efficiency of the network (Supporting Figure S3). Importantly, these two metrics capture distinct aspects of allosteric function: SPC reflects the structural position of a residue within the network, while Z-ASPL reports on the functional consequences of mutating that residue. The convergence of high SPC with high positive Z-ASPL at specific residues identifies the most critical allosteric hotspots—positions that are both structurally central and functionally sensitive to mutation. Conversely, residues with near-zero Z-ASPL values indicate positions where mutations do not significantly affect the average short path length, suggesting that these residues are not critical for maintaining global communication efficiency, regardless of their structural position.

**Figure 7.**
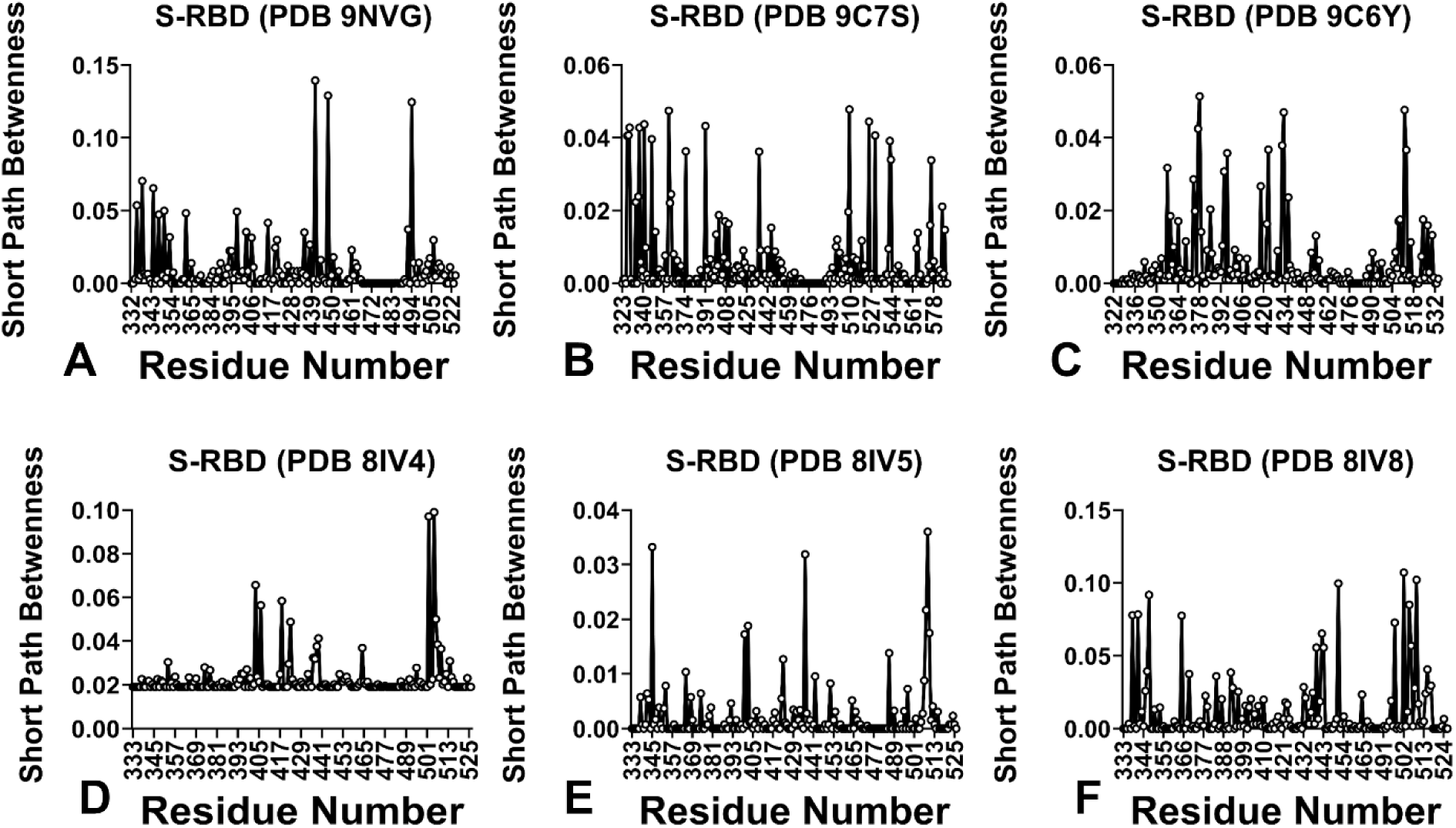
Allosteric network analysis: Short Path Betweenness (SPC) profiles. SPC profiles revealing communication hubs and mechanical pathways across the six antibody-RBD complexes. Higher SPC values indicate residues serving as critical conduits for communication between distant regions of the protein. (A) COV2-3835 (9NVG). (B) COV2-3891 (9C7S). (C) COV2-3906 (9C6Y). (D) 8IV5 complex (class 1 antibody 8H12 + class 3 antibody 1C4). (E) 8IV4 complex (class 1 antibody 8H12 + class 4 antibody 3E2). (F) 8IV8 complex (class 3 antibody 1C4 + class 4 antibody 3E2). The SPC profiles establish the β-sheet core as the critical allosteric conduit for class 4 antibodies, distinguishing them from class 3 antibodies that show only localized communication.

The SPC profiles for the class 3 antibody complexes (Figure 7A-B, 7D) reveal a pattern of localized communication. For COV2-3835 (Figure 7A), the SPC profile shows minimal elevated betweenness values, with only modest peaks within the β4-β5 hairpin region (ASN434, LYS441). Critically, the Z-ASPL values across the RBD remain near zero (Supporting Figure S3A), indicating that mutations at most positions do not substantially alter the global communication network. There are no positions where both SPC and Z-ASPL converge to identify strong allosteric hotspots—residues with elevated SPC do not exhibit correspondingly positive Z-ASPL values, suggesting that while certain residues serve as local structural hubs, their mutation does not disrupt the overall communication efficiency. This is consistent with neutralization through physical occlusion without requiring long-range communication.

For COV2-3891 (Figure 7B), the SPC profile shows modest peaks at VAL445 and GLY446 within the β4-β5 hairpin, with minimal values elsewhere. The class 1/class 3 dual complex (Figure 7D) shows elevated SPC at the RBM loop (PHE486, TYR489, GLN493) and β4-β5 hairpin (VAL445), but again, no strong convergence of SPC and Z-ASPL, and SPC values remain localized with no significant propagation through the β-sheet core. The Z-ASPL values in these complexes remain near zero for residues outside the immediate binding interface (Supporting Figure S3B, S3D), confirming that class 3 antibody engagement does not establish allosteric networks that connect the lateral face to distal functional regions. The absence of convergence between SPC and Z-ASPL in class 3 complexes indicates that the structural hubs identified by SPC do not correspond to functionally critical nodes whose mutation would disrupt allosteric communication.

In striking contrast, the SPC profiles for the class 4 antibody complexes (Figure 7C, 7E-F) reveal extensive allosteric communication networks spanning the entire RBD structure, and critically, these SPC peaks converge with positive Z-ASPL peaks at specific residues (Supporting Figure S3C, S3E-F). For the single class 4 antibody (Figure 7C), the SPC profile shows elevated betweenness throughout the β-sheet core, establishing a continuous conduit from the cryptic inner face to the RBM loop. The hydrophobic core residues (TYR369, PHE377, THR385) show elevated SPC and positive Z-ASPL values, indicating their dual role as communication hubs whose mutation would severely disrupt the network. Critically, GLY404, ASP405, and ARG408 represent the most significant allosteric hotspots, with the highest SPC values and most positive Z-ASPL peaks. These residues serve as the primary conduits for allosteric signal propagation, and the convergence of both metrics at these positions establishes them as essential nodes where both structural centrality and functional sensitivity coincide. ASN437 also shows elevated SPC and positive Z-ASPL values. The Z-ASPL values become progressively less positive from the β-sheet core to the RBM loop, indicating a hierarchical organization where mutations at the core communication hubs would most severely disrupt the network, while mutations at peripheral positions would have more moderate effects. SPC values remain elevated throughout the RBM loop (470-490), with Z-ASPL values that, while less positive than those in the β-sheet core, indicate that mutations at these positions would still disrupt the communication pathway. This gradient—from the most positive Z-ASPL at the β-sheet core to less positive at the RBM loop— reflects the hierarchical organization of the allosteric network and explains why escape mutations are more likely at peripheral positions than at core communication hubs.

The class 1/class 4 dual complex (Figure 7E) shows a more moderate communication network, with elevated SPC at the hydrophobic core (372-376) and β-sheet core (404-408, 437). ARG408 and ASP405 emerge as the most significant allosteric hotspots where SPC and Z-ASPL converge, though with less extreme Z-ASPL values than the single class 4 antibody. The RBM loop shows elevated SPC and Z-ASPL at residues such as PHE486 and TYR489, reflecting the balance between allosteric communication and direct constraint. The dual class 3/class 4 complex (Figure 7F) shows the most extensive communication network, with enhanced betweenness throughout the β-sheet core, residues GLY404, ASP405, ARG408, and ASN437 showing the highest SPC values observed. Z-ASPL values are most positive for these residues (Supporting Figure S3F), indicating mutations would most severely disrupt the amplified allosteric network. Critically, RBM loop residues show enhanced SPC with Z-ASPL values for TYR489, GLN493, and TYR505 more positive than in other class 4 complexes, indicating that dual engagement amplifies propagation to the receptor-binding surface and makes even peripheral RBM loop residues more critical for the communication network. The convergence of SPC and Z-ASPL at these positions—where both metrics identify the same residues as critical— provides compelling evidence that these residues are essential nodes whose structural position and chemical identity are both crucial for allosteric communication.

To visualize the spatial distribution of allosteric hotspots and their relationship to binding epitopes, we projected residues with high SPC and positive Z-ASPL values onto the RBD surface for representative class 3 (COV2-3835, 9NVG) and class 4 (COV2-3906, 9C6Y) antibody complexes (Figure 8). This structural projection reveals a striking difference in the organization of allosteric communication pathways between the two antibody classes. For the class 3 antibody complex (Figure 8A), residues with high SPC and positive Z-ASPL (red) are predominantly confined to the binding epitope (blue) and the immediate binding hotspots (orange). The allosteric hotspots do not extend beyond the lateral face interface, consistent with the localized nature of mechanical perturbation and the absence of long-range communication through the β-sheet core. The RBM loop region shows minimal allosteric hotspot distribution, confirming that class 3 antibody engagement does not establish communication pathways that connect the binding interface to distal functional regions. In striking contrast, for the class 4 antibody complex (Figure 8B), the distribution of allosteric hotspots (red) extends well beyond the binding epitope (blue) and binding hotspots (orange), forming a continuous spatial gradient that propagates from the cryptic inner face through the β-sheet core toward the ACE2-binding site. The allosteric hotspots are concentrated along the β-sheet core residues GLY404, ASP405, ARG408, and ASN437, which serve as the primary conduits for signal propagation.

**Figure 8.**
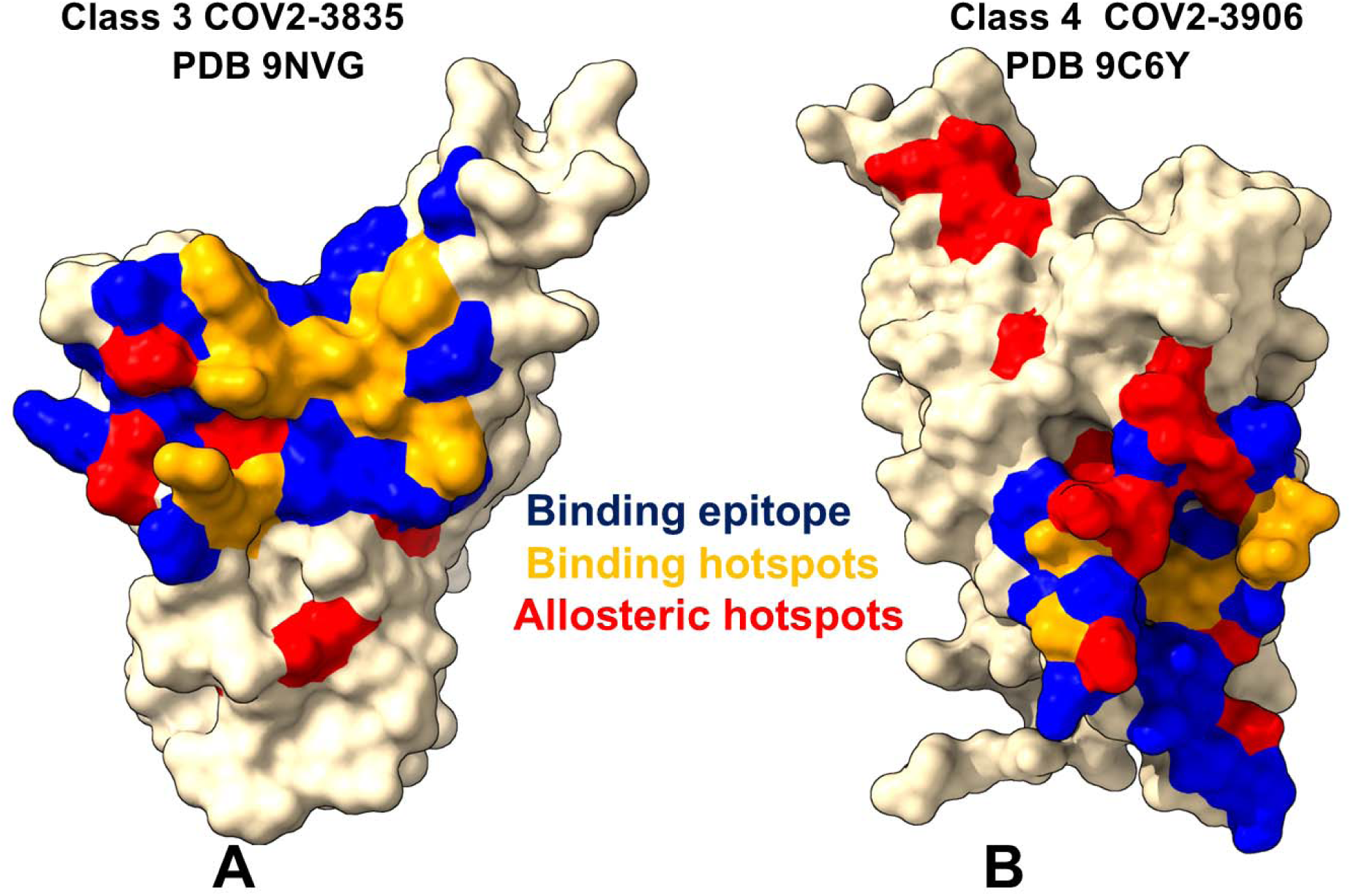
Spatial distribution of allosteric hotspots on the RBD surface. Projection of residues with high SPC and strongly negative Z-ASPL values (red) onto the RBD surface for representative class 3 (A, COV2-3835, PDB 9NVG) and class 4 (B, COV2-3906, PDB 9C6Y) antibody complexes. Binding epitope residues are shown in blue, binding hotspots (residues with highest mutational sensitivity) in orange. (A) Class 3 antibody complex showing allosteric hotspots confined to the binding epitope and immediate binding hotspots, with no propagation to the RBM loop or distal regions. (B) Class 4 antibody complex showing extensive allosteric hotspot distribution extending from the cryptic inner face through the β-sheet core toward the ACE2-binding site and RBM loop, establishing a continuous communication pathway. The spatial gradient of allosteric hotspots (red) from the binding interface through the β-sheet core to the RBM loop provides visual evidence for the long-range allosteric communication mechanism of class 4 antibodies. RBD is shown in surface representation (gray) with the ACE2-binding site indicated.

Critically, the allosteric hotspots extend into the RBM loop region (residues 470-490), establishing a communication pathway that connects the inner face to the receptor-binding surface. This spatial distribution provides direct structural evidence for the long-range allosteric communication mechanism of class 4 antibodies, demonstrating that engagement of the cryptic inner face induces conformational changes that propagate through the β-sheet core to the RBM loop. The gradient of allosteric hotspots—from the binding interface through the β-sheet core to the RBM loop—reflects the hierarchical organization of the allosteric network and explains why class 4 antibodies can induce allosteric loosening of the RBM loop despite binding at a distal site.

The network analysis thus identifies the β-sheet core as the critical structural conduit for allosteric communication in class 4 antibodies. The residues GLY404, ASP405, ARG408, and ASN437 consistently show the highest SPC values and most positive Z-ASPL values, establishing them as the primary communication hubs connecting the hydrophobic core anchor to the RBM loop. The convergence of both metrics at these positions—where high SPC indicates structural centrality and positive Z-ASPL indicates functional sensitivity to mutation—provides compelling evidence that these residues are essential nodes whose perturbation would severely compromise allosteric communication. This structural conduit is absent in class 3 antibody complexes, where the β-sheet core shows minimal SPC and near-zero Z-ASPL values, and where no convergence of the two metrics identifies critical allosteric hotspots. The spatial projection of allosteric hotspots (Figure 8) visually confirms that class 4 antibodies establish a long-range communication pathway that spans from the inner face to the receptor-binding surface, while class 3 antibodies induce only localized mechanical effects confined to the lateral face interface. The structural basis for this differential communication lies in the distinct epitope architectures: class 4 antibodies engage the hydrophobic core and β-sheet core through all six CDRs, establishing a mechanical pathway that propagates conformational changes to the RBM loop, while class 3 antibodies engage the lateral face without contacting the β-sheet core, limiting mechanical effects to the binding interface.

## Discussion

The integrated structural, dynamical, mechanical, energetic, and network analyses presented in this study establish a comprehensive biophysical framework that distinguishes the neutralization mechanisms of class 3 and class 4 antibodies. Our findings reveal that these two antibody classes achieve neutralization through fundamentally distinct strategies—localized mechanical constraint for class 3 antibodies versus long-range allosteric communication for class 4 antibodies—and that these mechanistic differences translate into markedly different mutational landscapes and escape vulnerabilities. Conformational dynamics analyses of class 3 antibody complexes consistently show that mechanical perturbation is strictly confined to the binding interface, with no significant propagation through the β-sheet core. The RBM loop exhibits moderate constraint, reflecting the antibody’s proximity to the receptor-binding surface, enabling effective steric hindrance without complete rigidification. The variation in RBM constraint across the class 3 antibody family correlates directly with epitope footprint extent—more extensive engagement, particularly of the α2-helix, leads to greater mechanical constraint. This establishes the mechanical state of the RBM loop as a key determinant of neutralization efficacy for class 3 antibodies. The mutational heatmaps reveal that class 3 epitopes exhibit a plastic periphery with a conserved β4-β5 hairpin anchor and variable sensitivity in the α2-helix and RBM loop, creating multiple escape pathways. The hierarchical organization of mutational sensitivity—primary hotspots at the β4-β5 hairpin, secondary hotspots at the α2-helix, and tertiary hotspots at the RBM loop—provides a quantitative basis for predicting which mutations are most likely to emerge under immune pressure. Importantly, our analysis focuses on RBD-level interactions, demonstrating that class 3 antibodies can impose mechanical constraint on the RBM loop through lateral face engagement.

The structural and dynamic diversity within the class 3 antibody family has important implications for understanding the differential neutralization potency and escape profiles of individual antibodies. COV2-3835, with its extensive engagement of the α2-helix, establishes a broader network of contacts that distributes mechanical constraint across multiple regions of the lateral face, resulting in greater RBM loop constraint and higher neutralization potency. In contrast, COV2-3891, with its compact footprint focused on the β4-β5 hairpin, imposes more limited mechanical constraint, correlating with reduced potency. This spectrum of engagement modes suggests that class 3 antibodies can be optimized for enhanced potency through engineering strategies that increase contact density and footprint extent on the lateral face. The class 1/class 3 dual complex demonstrates that combining a class 3 antibody with a class 1 antibody can enhance RBM loop constraint through dual steric blockade, achieving greater mechanical constraint than either antibody alone. This synergistic effect has important implications for antibody cocktail design, as combinations of class 3 and class 1 antibodies may achieve both high potency and broad protection.

In striking contrast, class 4 antibodies exhibit a dramatic mechanical dichotomy: extreme rigidification at the cryptic epitope coupled with pronounced allosteric loosening of the RBM loop. The network analysis identifies the β-sheet core residues as critical communication hubs. This explains why escape mutations are more likely at peripheral positions than at the core communication hubs—mutations at the β-sheet core would severely disrupt the allosteric network and compromise RBD structural integrity. The allosteric mechanism of class 4 antibodies represents a fundamentally different strategy for achieving neutralization. Rather than directly blocking ACE2 binding through steric occlusion, class 4 antibodies exploit the intrinsic mechanical connectivity of the RBD, using the β-sheet core as a conduit to transmit conformational changes from the inner face to the RBM loop. This indirect mechanism, while less potent than direct steric blockade, offers the advantage of targeting a highly conserved epitope that is resistant to mutational escape. The magnitude of the allosteric effect scales with the extent of inner-face engagement—COV2-3906, with its most extensive engagement through all six CDRs, shows the strongest mechanical gradient and RBM loosening. The class 1/class 4 dual complex shows a more moderate effect, reflecting 3E2’s focused footprint. The dual class 3/class 4 complex demonstrates that cooperative engagement of the inner face and lateral face can amplify the allosteric communication, producing the most extreme RBM loosening observed. This amplification arises from the combined anchoring effect—3E2 immobilizes the hydrophobic core while 1C4 provides additional structural stabilization through lateral face engagement—creating a more extensive footprint that enhances propagation of mechanical changes through the β-sheet core.

The convergence of our computational predictions with experimental DMS data provides robust validation of our approach.^42–45,118,119^ For class 3 antibodies, our predicted hotspots at the β4-β5 hairpin (LYS437, VAL445, GLY446) and α2-helix (ARG343, THR342) correspond to experimentally validated escape residues, including R346T, K444T, and L452R mutations that have emerged in circulating variants. The RBM loop hotspots (PHE486, TYR489) correspond to residues under dual selective pressure—mutations that escape antibody binding may also impair ACE2 binding, limiting their fitness. The agreement between our predicted hotspots and DMS escape profiles confirms that mutational sensitivity reflects underlying energetic importance, with VDW-driven hydrophobic packing as the primary determinant of binding affinity. For class 4 antibodies, our analysis identifies the hydrophobic core (TYR369, PHE377, THR385) and β-sheet core as the most critical hotspots, consistent with DMS studies showing these positions are highly conserved with escape mutations rarely observed due to prohibitive fitness costs. The main-chain electrostatic interactions we identified involving N370, K378, S383, and T385 provide a mechanistic explanation for this conservation—these interactions depend on backbone conformation rather than side-chain identity, and mutations disrupting them would require structurally disfavored backbone rearrangements.

The mechanistic principles identified in this study are consistent with the broader literature on class 3 and class 4 antibodies. Class 3 antibodies, including S309 (sotrovimab) and LY-CoV555, have been shown to target the lateral face and mediate neutralization through steric hindrance.^40,41,118,119^ The variation in epitope footprint and neutralization potency among class 3 antibodies reflects the structural plasticity of the lateral face, which can be engaged through multiple binding modes with different mechanical consequences. The conserved β4-β5 hairpin anchor identified in our analysis provides a stable contact point that is targeted across the class 3 antibody family, explaining why this region remains a consistent target despite variation in other parts of the epitope. Class 4 antibodies, including CR3022 and S304, have been shown to target the cryptic inner face and mediate neutralization through allosteric mechanisms.^40,41,118,119^ The extreme conservation of the hydrophobic core and β-sheet core across sarbecoviruses explains the ultra-broad binding of class 4 antibodies, while the indirect allosteric mechanism explains their limited neutralization potency. The identification of the β-sheet core as a critical allosteric conduit provides a mechanistic explanation for how class 4 antibodies achieve long-range communication from the inner face to the RBM loop.

The mutational landscapes of class 3 and class 4 antibodies reveal complementary vulnerabilities that have shaped the evolutionary trajectory of SARS-CoV-2. Class 3 antibodies face moderate escape vulnerability, with mutations concentrated in the α2-helix and RBM loop regions where mutational tolerance is higher. The recurrence of mutations such as R346T, K444T, and L452R across circulating variants reflects the selective pressure exerted by class 3 antibodies on these accessible regions of the lateral face. These mutations achieve immune escape without compromising RBD stability or ACE2 binding affinity, making them evolutionarily favorable. In contrast, class 4 antibodies maintain ultra-broad binding due to the immutable nature of the hydrophobic core and β-sheet core. The extreme sensitivity of these residues to mutation combined with their essential role in maintaining RBD structural integrity creates a high genetic barrier to resistance. This explains why escape mutations at the hydrophobic core are rarely observed in circulating variants, and why class 4 antibodies maintain efficacy across diverse lineages. The complementary nature of these mutational landscapes suggests that antibody cocktails combining class 3 and class 4 antibodies could provide both high potency and broad protection, limiting the emergence of escape variants. The integrated analysis provides a quantitative framework for guiding antibody design and predicting resistance. For class 3 antibodies, engineering strategies should focus on distributing binding energy across multiple hotspots to enhance resilience. Antibodies that distribute binding energy across multiple hotspots are likely to be more resilient to escape than those that concentrate energy at a single anchor as mutations that disrupt one hotspot can be compensated by persistent contacts at others. The dual class 3/class 4 complex demonstrates that cooperative engagement of the inner face and lateral face can amplify allosteric communication, suggesting that engineering strategies combining class 3 and class 4 antibodies could achieve both high potency and broad protection.

### Conclusions

This study unveils the dynamic and mechanistic dichotomy between class 3 and class 4 antibodies through a multi-layered biophysical analysis. The findings establish that epitope location on the RBD determines not only binding geometry but also the physical strategy by which neutralization is achieved. Class 3 antibodies impose localized mechanical constraint on the RBM loop, with perturbation strictly confined to the lateral face interface. Class 4 antibodies, by contrast, establish a long-range allosteric conduit that transmits mechanical changes from the hydrophobic core through the β-sheet core to induce pronounced RBM loosening. This distinction explains why class 3 antibodies achieve higher potency through direct mechanical interference with the receptor-binding surface, while class 4 antibodies achieve ultra-broad binding through targeting an evolutionarily constrained structural core.

The mutational landscapes of the two classes reveal a fundamental asymmetry in viral escape potential. Class 4 epitopes are anchored by an immutable hydrophobic core whose residues are exquisitely sensitive to mutation yet invariant across sarbecoviruses due to their essential role in RBD structural integrity. The main-chain electrostatic interactions at the 370s loop and β-sheet core further reinforce this barrier, as they depend on backbone conformation rather than side-chain identity. Class 3 epitopes exhibit a plastic periphery with a conserved β4-β5 hairpin anchor and variable sensitivity in the α2-helix and RBM loop, creating a hierarchical organization of escape pathways. This asymmetry explains the differential evolutionary pressures on these antibody classes: class 3 antibodies are subject to ongoing immune escape through mutations at accessible lateral face positions, while class 4 antibodies maintain efficacy against diverse variants due to the high fitness cost of mutations at the hydrophobic core.

The allosteric network analysis identifies the β-sheet core residues GLY404, ASP405, ARG408, and ASN437 as essential communication hubs where structural centrality and functional sensitivity converge. The continuous spatial gradient of allosteric hotspots extending from the cryptic inner face through the β-sheet core to the RBM loop provides direct evidence for the long-range communication pathway exploited by class 4 antibodies. This pathway is entirely absent in class 3 antibody complexes, confirming that their neutralization mechanism does not rely on allosteric communication. The identification of these evolutionarily constrained allosteric conduits as vulnerable targets offers a promising strategy for developing antibodies with durable efficacy against rapidly evolving pathogens.

This integrated analysis provides a quantitative foundation for guiding antibody design. For class 3 antibodies, distributing binding energy across multiple hotspots enhances resilience against escape, as mutations disrupting one hotspot can be compensated by persistent contacts at others. For class 4 antibodies, engaging multiple regions of the inner face could improve potency while maintaining the high genetic barrier to resistance. The complementary mutational landscapes and mechanical strategies of the two classes suggest that antibody cocktails combining both could provide both high potency and broad protection. The computational framework developed here provides a generalizable approach for understanding antibody neutralization mechanisms and predicting immune escape across diverse viral targets, with implications for the rational design of next-generation antibody therapeutics that balance potency, breadth, and resilience.

## Conflicts of Interest

The authors declare no conflict of interest. The funders had no role in the design of the study; in the collection, analyses, or interpretation of data; in the writing of the manuscript; or in the decision to publish the results.

## Author Contributions

Conceptualization, G.V.; methodology, G.V.; software, M.A., W.G., B.F., M.L., L.T., G.V.; validation, M.A., W.G., B.F., M.L., L.T., G.V.; formal analysis, M.A., W.G., B.F., M.L., L.T., G.V.; investigation, M.A., W.G., B.F., M.L., L.T., G.V.; resources, M.A., W.G., B.F., M.L., L.T., G.V.; data curation, M.A., W.G., B.F., M.L., L.T., G.V.; writing— original draft preparation, M.A., W.G., B.F., M.L., L.T., G.V.; writing—review and editing, G.V.; visualization, M.A., W.G., B.F., M.L., L.T., G.V.; supervision, G.V.; project administration, G.V.; funding acquisition, G.V. All authors have read and agreed to the published version of the manuscript.

## Funding

This research was funded by the National Institutes of Health under Award 1R01AI181600-01, 5R01AI181600-02 and Subaward 6069-SC24-11 to G.V.

## Data Availability Statement

Data is fully contained within the article and Supplementary Materials. Crystal structures were obtained and downloaded from the Protein Data Bank (http://www.rcsb.org). The rendering of protein structures was done with UCSF ChimeraX package (https://www.rbvi.ucsf.edu/chimerax/) and Pymol (https://pymol.org/2/). All mutational heatmaps were produced using the developed software that is freely available at https://alshahrani.shinyapps.io/HeatMapViewerApp/.

## Acknowledgments

The authors acknowledge support from Schmid College of Science and Technology at Chapman University for providing computing resources at the Keck Center for Science and Engineering.

## Notes

### Competing Interest Statement

The authors have declared no competing interest.

