## Supplementary Figures S1-S3 for "Localized Rigidification and Allosteric Modulation Mechanisms of SARS-CoV-2 Spike Neutralization by Class 3 and Class 4 Antibodies at Atomic Resolution: An Integrated Computational Study of Binding, Dynamics, and Allostery"

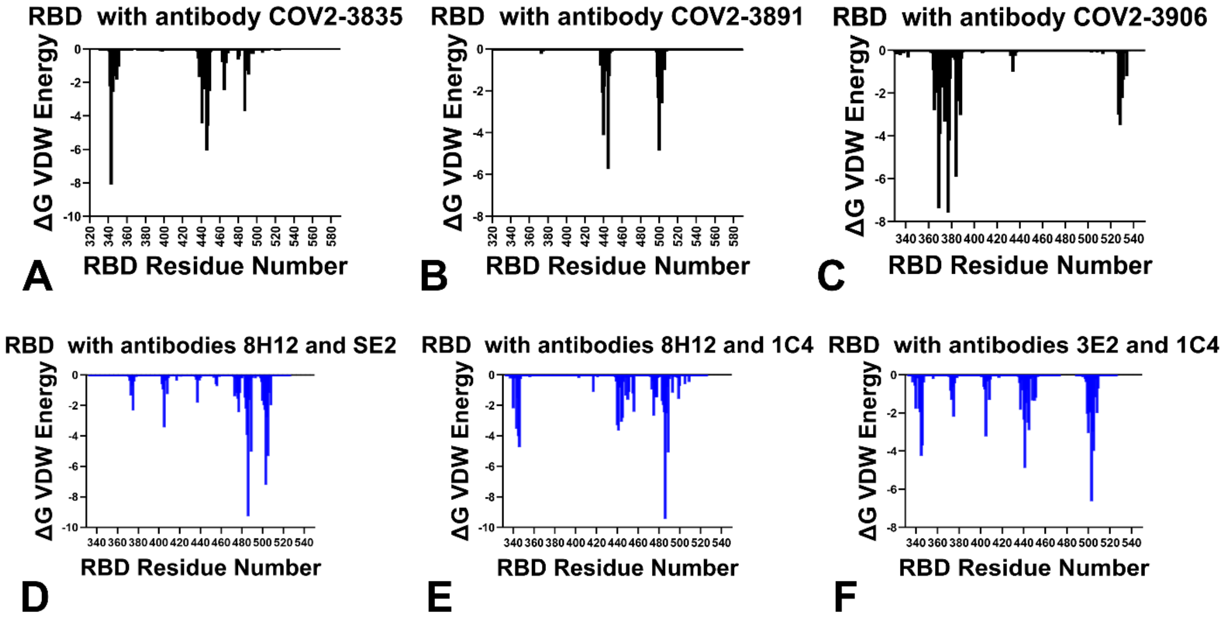


**Figure S1. Van der Waals energy contribution profiles from MM-GBSA analysis.** Per-residue van der Waals (VDW) binding energy contributions across the six antibody-RBD complexes. Negative values indicate favorable van der Waals interactions. (A) COV2-3835 (9NVG). (B) COV2-3891 (9C7S). (C) COV2-3906 (9C6Y). (D) Class 1/class 4 dual complex (8IV4). (E) Class 1/class 3 dual complex (8IV5). (F) Class 3/class 4 dual complex (8IV8). VDW contributions serve as the dominant driver of binding for class 3 antibodies, with energies concentrated at the α2-helix, β4-β5 hairpin, and RBM loop. For class 4 antibodies, VDW contributions reveal extreme stabilization at the hydrophobic core, explaining the catastrophic effect of mutations at these positions.

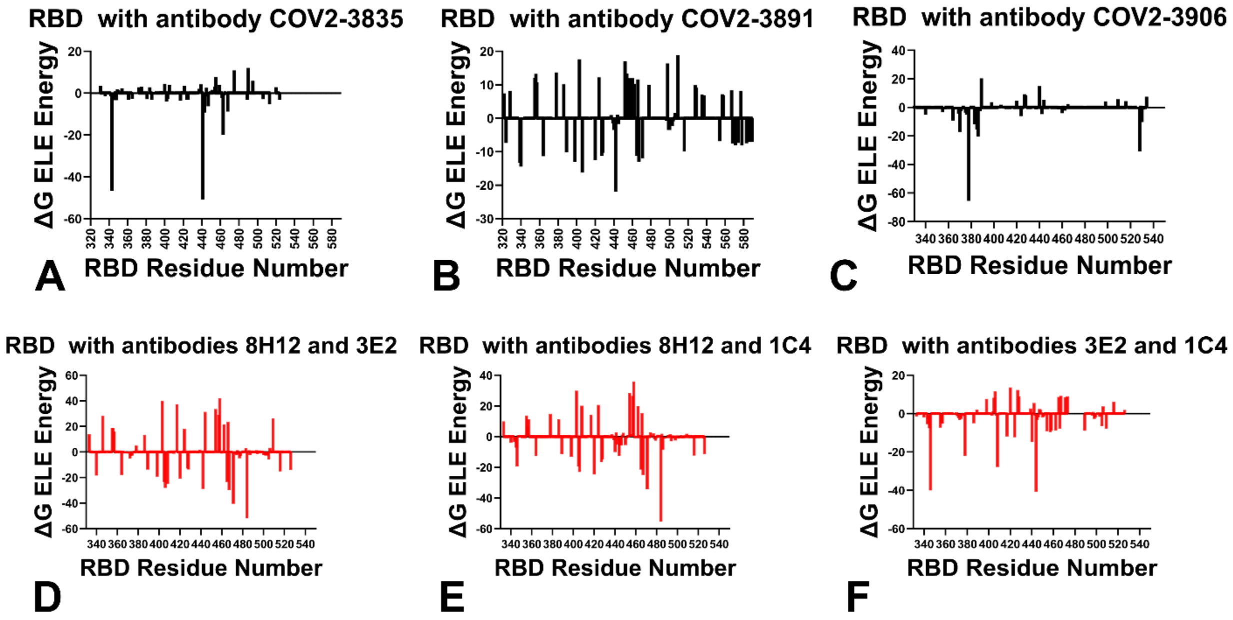


**Figure S2. Electrostatic energy contribution profiles from MM-GBSA analysis.** Per-residue electrostatic (ELE) binding energy contributions across the six antibody-RBD complexes. Negative values indicate favorable electrostatic interactions. (A) COV2-3835 (9NVG). (B) COV2-3891 (9C7S). (C) COV2-3906 (9C6Y). (D) Class 1/class 4 dual complex (8IV4). (E) Class 1/class 3 dual complex (8IV5). (F) Class 3/class 4 dual complex (8IV8). Electrostatic contributions play a secondary role for class 3 antibodies, with ARG343 showing the most favorable interactions partially offset by solvation. For class 4 antibodies, electrostatic contributions reveal extensive main-chain hydrogen bonds involving the 370s loop (N370, K378, S383, T385) and β-sheet core, which are resistant to mutation as they depend on backbone conformation rather than side-chain identity.

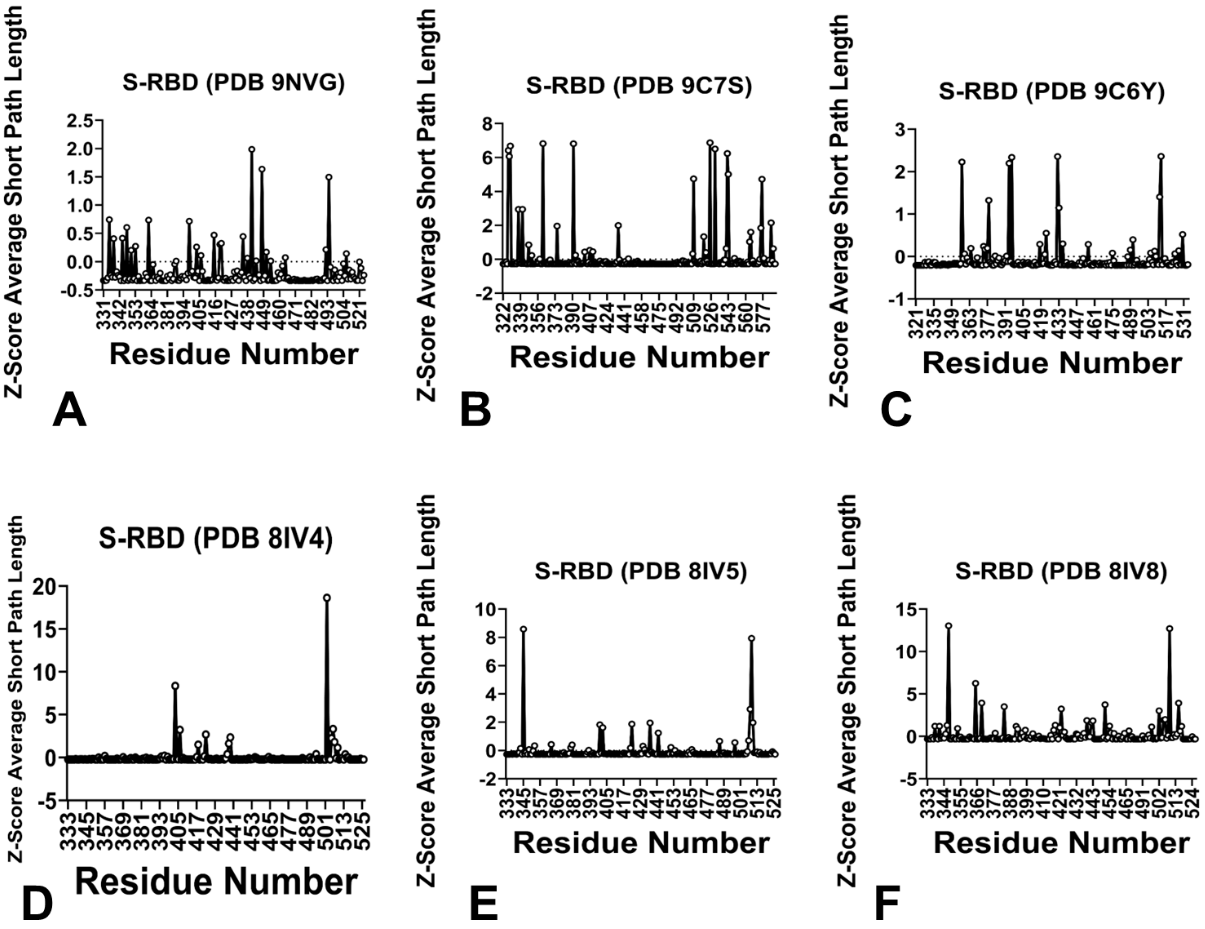


**Figure S3. Z-score of Average Short Path Length (Z-RCA) profiles from allosteric network analysis.** Z-RCA profiles identifying residues whose mutational perturbation significantly alters the global communication efficiency of the protein structure network. More negative Z-RCA values indicate that mutations at these positions would most severely disrupt the allosteric communication network. (A) COV2-3835 (9NVG). (B) COV2-3891 (9C7S). (C) COV2-3906 (9C6Y). (D) Class 1/class 3 dual complex (8IV5). (E) Class 1/class 4 dual complex (8IV4). (F) Class 3/class 4 dual complex (8IV8). The convergence of high SPC values (Figure 7) with strongly negative Z-RCA values identifies the most critical allosteric hotspots. For class 4 antibodies, the β-sheet core residues GLY404, ASP405, ARG408, and ASN437 show the most negative Z-RCA values, establishing them as essential communication hubs connecting the hydrophobic core to the RBM loop. In contrast, class 3 antibody complexes show minimal Z-RCA values, consistent with localized mechanical perturbation and the absence of long-range allosteric communication.
